# Shared genetic background between children and adults with attention deficit/hyperactivity disorder

**DOI:** 10.1101/589614

**Authors:** Paula Rovira, Ditte Demontis, Cristina Sánchez-Mora, Tetyana Zayats, Marieke Klein, Nina Roth Mota, Heike Weber, Iris Garcia-Martínez, Mireia Pagerols, Laura Vilar, Lorena Arribas, Vanesa Richarte, Montserrat Corrales, Christian Fadeuilhe, Rosa Bosch, Gemma Español Martin, Peter Almos, Alysa E. Doyle, Eugenio Horacio Grevet, Oliver Grimm, Anne Halmøy, Martine Hoogman, Mara Hutz, Christian P. Jacob, Sarah Kittel-Schneider, Per M. Knappskog, Astri J. Lundervold, Olga Rivero, Diego Luiz Rovaris, Angelica Salatino-Oliveira, Bruna Santos da Silva, Evgenij Svirin, Emma Sprooten, Tatyana Strekalova, ADHD Working Group of the Psychiatric Genomics Consortium, 23andMe Research team, Alejandro Arias-Vasquez, Edmund J.S. Sonuga-Barke, Philip Asherson, Claiton Henrique Dotto Bau, Jan K. Buitelaar, Bru Cormand, Stephen V. Faraone, Jan Haavik, Stefan E. Johansson, Jonna Kuntsi, Henrik Larsson, Klaus-Peter Lesch, Andreas Reif, Luis Augusto Rohde, Miquel Casas, Anders D. Børglum, Barbara Franke, Josep Antoni Ramos-Quiroga, María Soler Artigas, Marta Ribasés

**Author notes:** These authors contributed equally to this work (Paula Rovira, Ditte Demontis, Anders D. Børglum, Barbara Franke, Josep Antoni Ramos-Quiroga, María Soler Artigas, Marta Ribasés). Correspondence and requests for materials should be addressed to M.R.) or M.S.A).

## Abstract

Attention deficit/hyperactivity disorder (ADHD) is a common neurodevelopmental disorder characterized by age-inappropriate symptoms of inattention, impulsivity and hyperactivity that persist into adulthood in the majority of the diagnosed children. Despite several risk factors during childhood predicting the persistence of ADHD symptoms into adulthood, the genetic architecture underlying the trajectory of ADHD over time is still unclear. We set out to study the contribution of common genetic variants to the risk for ADHD across the lifespan by conducting meta-analyses of genome-wide association studies on persistent ADHD in adults and ADHD in childhood separately and comparing the genetic background between them in a total sample of 17,149 cases and 32,411 controls. Our results show nine new independent loci and support a shared contribution of common genetic variants to ADHD in children and adults. No subgroup heterogeneity was observed among children, while this group consists of future remitting and persistent individuals. We report similar patterns of genetic correlation of ADHD with other ADHD-related datasets and different traits and disorders among adults, children and when combining both groups. These findings confirm that persistent ADHD in adults is a neurodevelopmental disorder and extend the existing hypothesis of a shared genetic architecture underlying ADHD and different traits to a lifespan perspective.

---

Attention deficit/hyperactivity disorder (ADHD) is a common neurodevelopmental disorder that severely impairs the daily functioning of patients due to age-inappropriate levels of impulsivity and hyperactivity, and/or difficulties in focusing attention. ADHD has a prevalence of 3.4% in childhood, and impairing symptoms persist into adulthood in around two-thirds of children with ADHD diagnosis, with an estimated adult prevalence around 3%^1,2^.

ADHD is a multifactorial disorder with heritability averaging 76% throughout the lifespan^3-5^. There is consistent evidence that both common and rare variants make an important contribution to the risk for the disorder^6-11^. Several genome-wide association studies (GWAS) and meta-analyses across those have been conducted^7^, but only the largest GWAS meta-analysis (GWAS-MA) performed to date, including 20,183 ADHD patients and 35,191 controls, reported genome-wide significant loci (N=12)^6^. This study concluded that common genetic variants (minor allele frequency, MAF, >0.01) account for 22% of the heritability of the disorder^6^ and supported substantial genetic overlap between ADHD and other brain disorders and behavioral/cognitive traits^12^.

The presentation of ADHD symptoms changes from childhood to adulthood, with lower levels of hyperactivity in adulthood but a high risk for ongoing attention problems, disorganization, and emotional dysregulation^13,14^. As in the general population, the pattern of psychiatric and somatic comorbid conditions in ADHD also changes substantially over time, with learning disabilities, oppositional defiant disorder, enuresis and conduct disorder being more prevalent in children, and substance use disorders, social phobia, insomnia, obesity, and mood disorders becoming more pronounced in adulthood^1,15-18^. In addition, persistent ADHD in adults is, compared to the general population (and to cases with remitting ADHD), associated with higher risk for a wide range of functional and social impairments, including unemployment, accidents and criminal behavior^7,19-23^

Several risk factors measured in childhood predict the persistence of ADHD symptoms into adulthood, such as the presence of comorbid disorders, the severity of ADHD symptoms, being exposed to psychosocial adversity as well as having a high polygenic risk score for childhood ADHD^24-28^. Twin studies suggest that both stable and dynamic genetic influences affect the persistence of ADHD symptoms^4,5,29,30^. However, specific genetic factors differentiating childhood and persistent ADHD into adulthood are not well understood due to the lack of longitudinal studies. Since molecular gene-finding studies, including the most recent GWAS-MA of ADHD, have been performed in children and adults either separately or jointly^6,31-40^, large-scale analyses comparing the genetic basis of children and adults with persistent ADHD are yet to be conducted.

Given this background, we set out to study the contribution of common genetic variants to the risk for ADHD from a lifespan perspective. We report, for the first time, a GWAS-MA on persistent ADHD in adults (according to DSM-IV/ICD-10 criteria), a GWAS-MA on ADHD in childhood (that may include remittent and persistent forms of the disorder), and the comparison of their genetic background.

## Results

### GWAS meta-analysis of persistent ADHD in adults

The GWAS-MA of persistent ADHD in adults included six datasets from the International Multi-centre persistent ADHD CollaboraTion (IMpACT) consortium, two datasets from the Psychiatric Genomics Consortium (PGC) and the subset of adults from the Lundbeck Foundation Initiative for Integrative Psychiatric Research (iPSYCH) cohort included in Demontis and Walters et al.^6^ (detailed information can be found in Supplementary Table 1 and in the Supplementary Material). In total, 22,406 individuals (6,532 adult ADHD cases and 15,874 controls) were included. The overall lambda (λ) value was 1.09 (λ_1000_ = 1.01) and the linkage disequilibrium (LD) score regression ratio was 0.13, suggesting minimal population stratification or other systematic biases (Supplementary Figure 1A). The proportion of heritability of persistent ADHD attributable to common single nucleotide polymorphisms (SNP-h^2^) was 0.21 (SE=0.026), with a nominally significant enrichment in the heritability of variants located in conserved genomic regions (P=5.18E-03) and in the cell-specific histone mark H3K4me1 (P=3.17E-02) (Supplementary Figure 2A). The gene-based analysis revealed six genes in four loci (*ST3GAL3, FRAT1/FRAT2, CGB1* and *RNF225/ZNF584*) significantly associated with persistent ADHD, with *ST3GAL3* being the most significant one (P=8.72E-07) (Supplementary Table 2A). The single-marker analysis showed no variants exceeding genome-wide significance (P<5.00E-08), with the most significant signal being rs3923931 (P=1.69E-07) (Figure 1A and Supplementary Table 3A). Similarly, no significant gene sets were identified in the pathway analysis after correction for multiple comparisons (Supplementary Table 4A [in excel]).

**Table 1.**
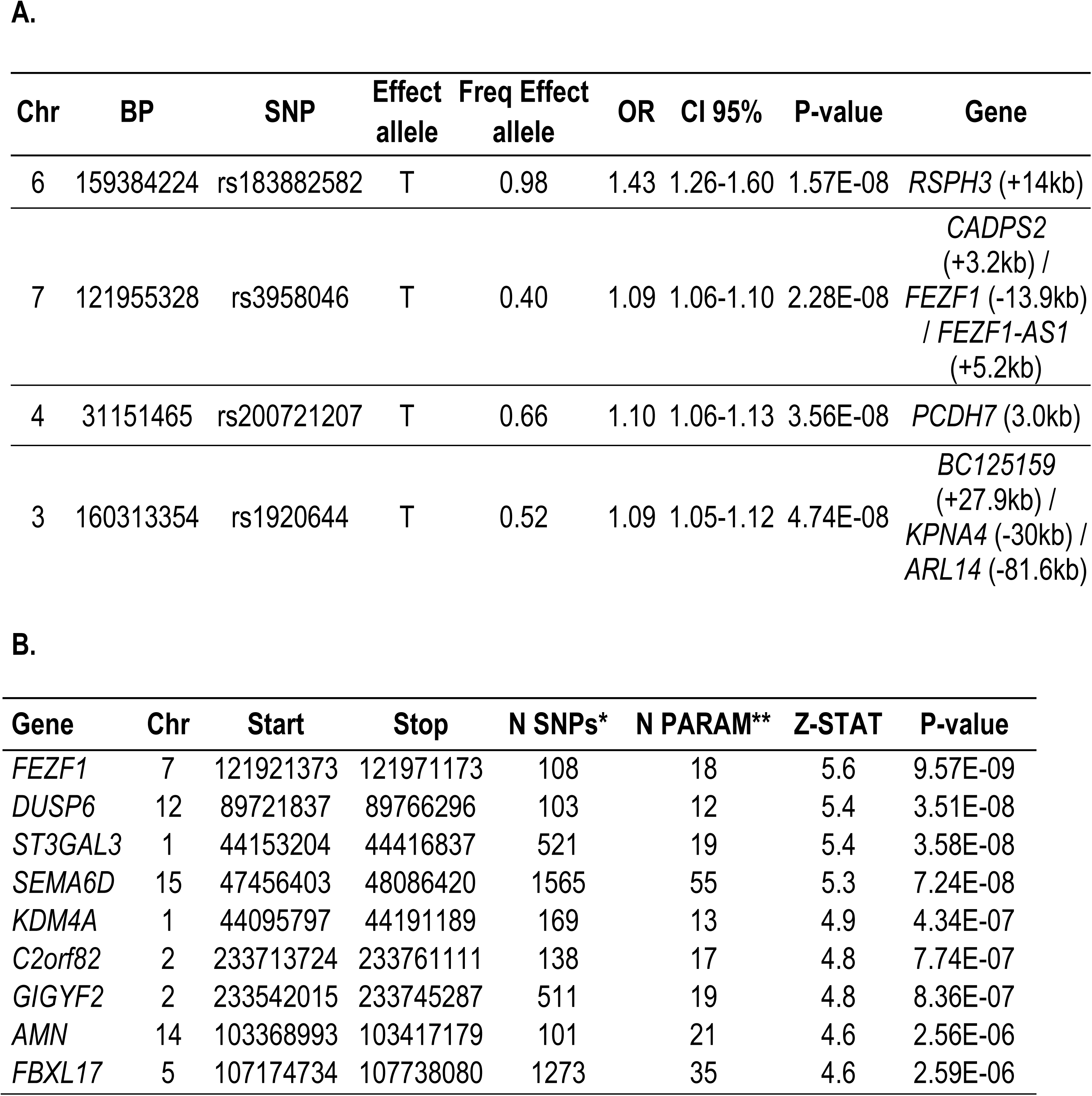
Genome-wide significant loci in the GWAS meta-analysis of ADHD across the lifespan identified through (A) single-variant analysis and (B) gene-based analysis. The location (chromosome (Chr) and base position (BP)), effect allele and its frequency, odds ratio (OR) of the effect allele with 95% confidence interval (CI 95%) and association P-values, along with genes in the locus are shown for each index variant ID (SNP). For the gene-based results, the number of single nucleotide polymorphisms in the genes (*) and the number of relevant parameters used in the model by MAGMA software^70^ (**) are given.

**Figure 1.**
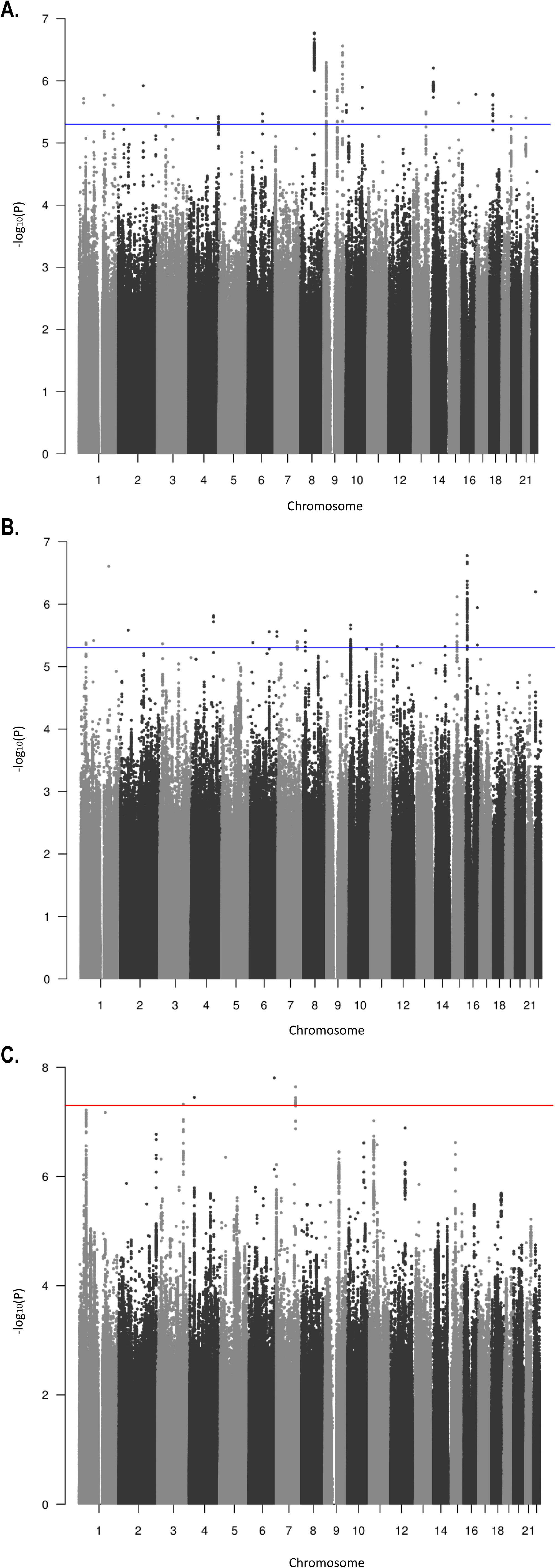
Manhattan plots of GWAS meta-analyses of (A) Nine cohorts of persistent ADHD in adults, (B) 10 cohorts of ADHD in childhood and (C) GWAS datasets of ADHD across the lifespan (ADHD in childhood + persistent ADHD). Horizontal lines indicate suggestive (P-value=5.00E-06) and genome-wide significant (P=5.00E-08) thresholds in A-B and C, respectively.

**Figure 2.**
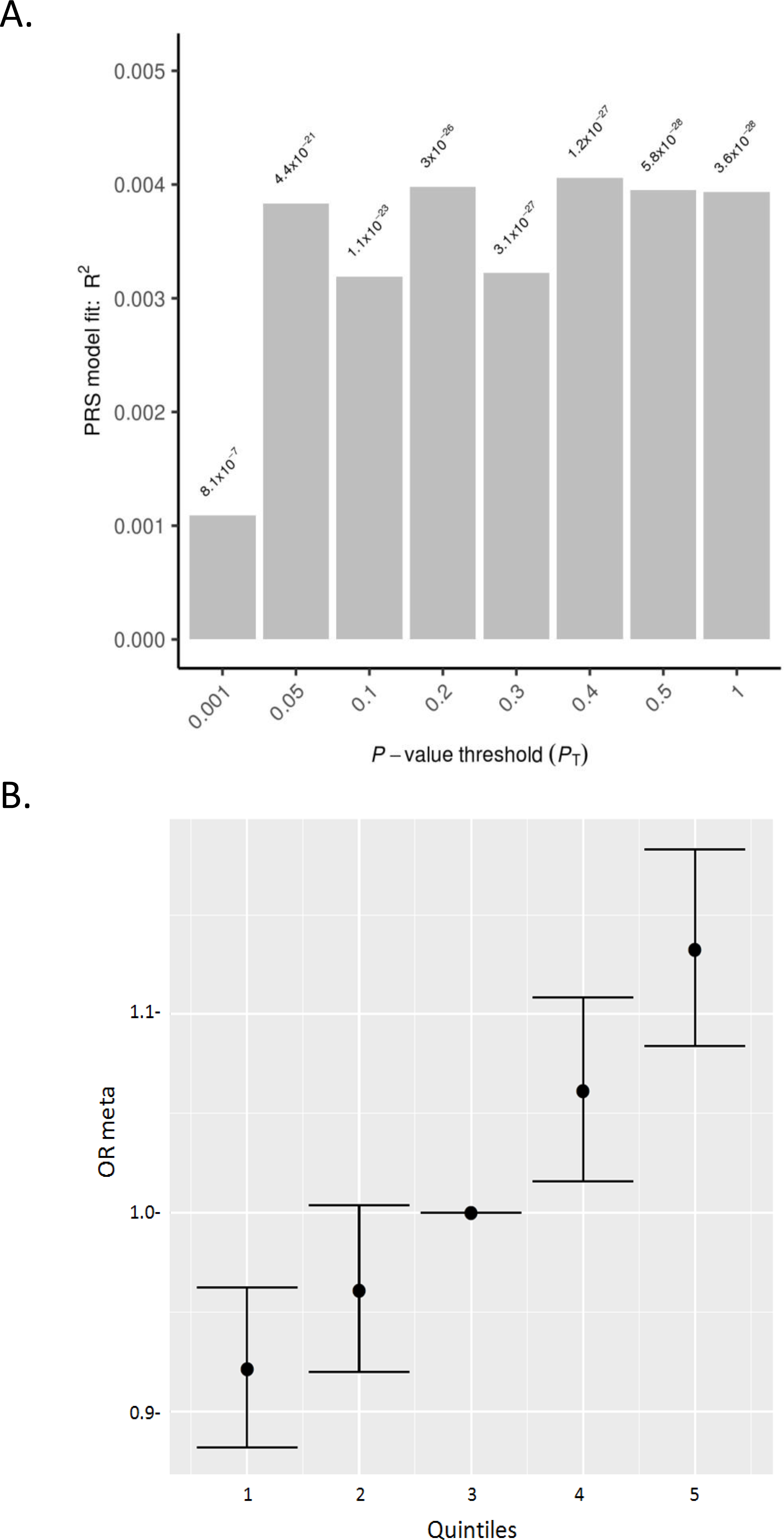
Polygenic risk scores for ADHD in childhood tested on persistent ADHD as target sample. (A) Bar plot and (B) Quintile plot of meta-analysis odds ratios (OR meta) with 95% confidence intervals for P-value threshold=0.4 using the third quintile as baseline.

### GWAS meta-analysis of ADHD in childhood

To compare the genetic background between persistent ADHD in adults and ADHD in childhood (that may include future remittent and persistent forms of the disorder), we conducted a GWAS-MA on children with ADHD in a total of 27,154 individuals (10,617 ADHD cases and 16,537 controls). The sample consisted of two datasets from Brazil and Spain, seven datasets from the PGC and the subset of children from the iPSYCH cohort included in Demontis and Walters et al.^6^ (detailed information can be found in Supplementary Table 1 and in the Supplementary Material). We found no evidence of genomic inflation or population stratification (λ=1.12 and λ_1000_ =1.01, LD score regression ratio=0.13) (Supplementary Figure 1B). The SNP-h^2^ for ADHD in childhood was 0.20 (SE=0.023), with a significant enrichment in the heritability of variants located in conserved genomic regions after Bonferroni correction (P=1.21E-06) (Supplementary Figure 2B). The gene-based analysis highlighted a significant association between *FEZF1* and ADHD in childhood (P=5.42E-07) (Supplementary Table 2B). No single genetic variant exceeded genome-wide significance (P<5.00E-08), with the top signal being in rs55686778 (P=1.67E-07) (Figure 1B and Supplementary Table 3B), and no significant gene sets were identified in the pathway analysis after correction for multiple comparisons (Supplementary Table 4B [in excel]).

### Comparison of the genetic background of persistent ADHD in adults and ADHD in childhood

We found a strong and significant genetic correlation between persistent ADHD in adults and ADHD in childhood (rg=0.81, SE=0.09, P=2.13E-21). Sign test results provided evidence of a consistent direction of effect of genetic variants associated with ADHD in children in persistent ADHD and vice-versa (P=6.60E-04 and P=4.47E-03, respectively for variants with P<5.00E-05 in either dataset) (Supplementary Table 5). In addition, Polygenic Risk Score (PRS) analyses showed that childhood ADHD PRSs were associated with persistent ADHD at different predefined P-value thresholds, with the P=0.40 threshold explaining the most variance (R^2^=0.0041 and P=1.20E-27) (Figure 2A). The quintiles of the PRS built using this threshold showed the expected trend of higher ADHD risk for individuals in higher quintiles (Figure 2B, Supplementary Table 6).

We tested whether the genetic correlation between persistent ADHD and ADHD in childhood was driven by a subset of children enriched for persistent ADHD-associated alleles using the Breaking Up Heterogeneous Mixture Based On Cross-locus correlations (BUHMBOX) analysis. We found no evidence of genetic heterogeneity in children, supporting that the sharing of persistent ADHD-associated alleles between children and adults was driven by the whole group of children, with a statistical power >98% and assuming 65% persistence (Supplementary Table 7).

### Meta-analysis of GWAS on ADHD across the lifespan

Given the strong genetic correlation between persistent ADHD in adults and ADHD in children, we performed a GWAS-MA of ADHD across the lifespan including all datasets available (nine GWAS of persistent ADHD in adults and 10 GWAS of ADHD in childhood). In total, we included 49,560 individuals (17,149 ADHD cases and 32,411 controls), and no evidence of genomic inflation or population stratification was found (λ=1.18 and λ_1000_ =1.01, LD score regression ratio=0.14) (Supplementary Figure 1C). The SNP-h^2^ for ADHD across the lifespan was 0.19 (SE=0.01), and a significant enrichment in the heritability of variants located in conserved genomic regions was observed after Bonferroni correction (P=1.53E-06) (Supplementary Figure 2C). We identified four genome-wide significant variants (Figure 1C, Figure 3, Table 1A and Supplementary Figure 3) and nine genes in seven loci (*FEZF1, DUSP6, ST3GAL3/KDM4A, SEMA6D, C2orf82/GIGYF2, AMN* and *FBXL17*) significantly associated with ADHD across the lifespan (Table 1B). The most significantly associated locus was on chromosome 6 (index variant rs183882582-T, OR=1.43 (95% CI 1.26-1.60), P=1.57E-08), followed by loci on chromosome 7 (index variant rs3958046), chromosome 4 (index variant rs200721207) and chromosome 3 (index variant rs1920644) (Table 1A, Figure 3). The gene-set analysis showed a significant association of the “ribonucleoprotein complex” GO term with ADHD across the lifespan (P.adj=0.021) (Supplementary Table 4C [in excel]).

**Figure 3.**
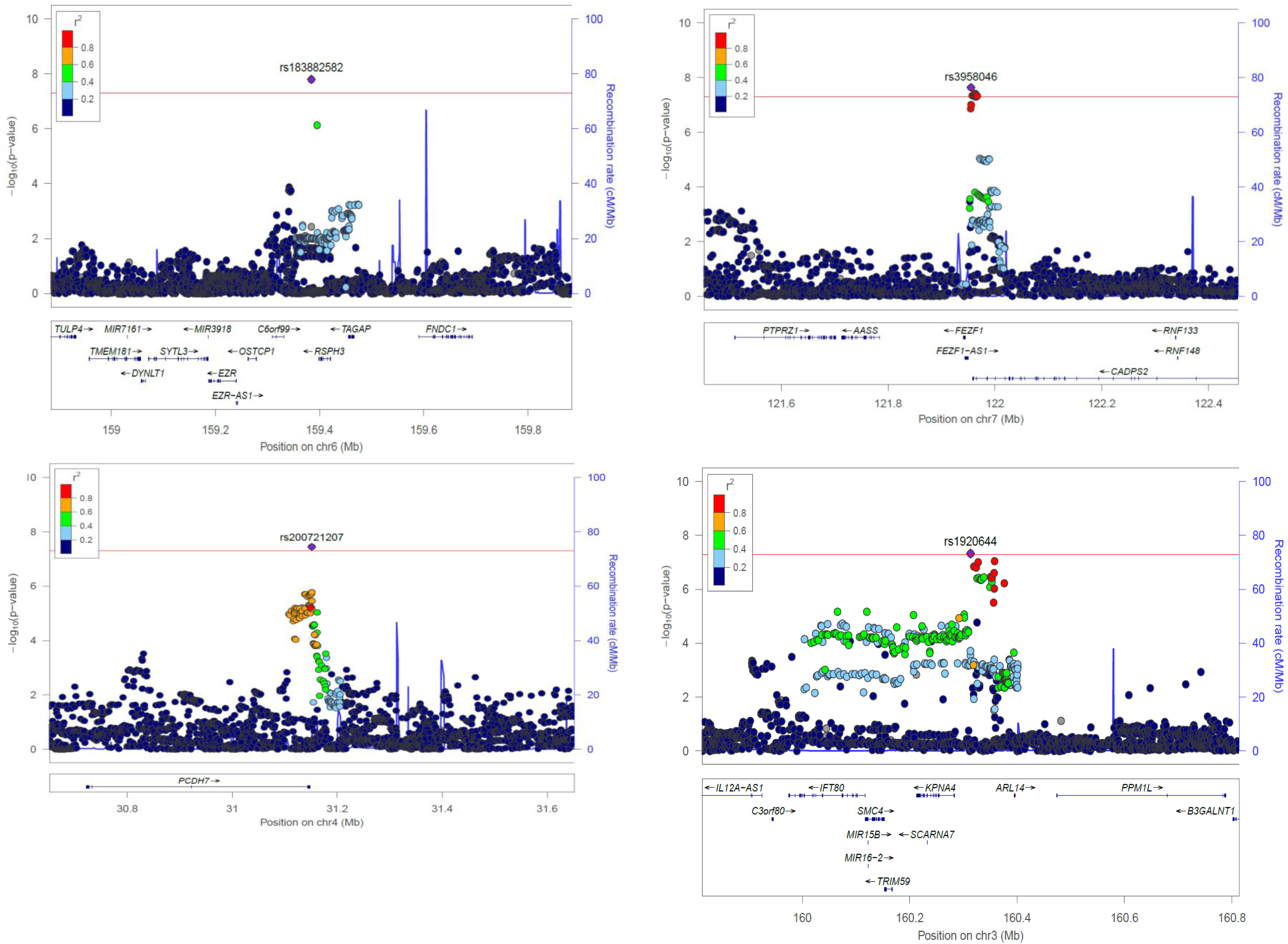
Regional association plots for genome-wide significant loci identified in the GWAS meta-analysis of ADHD across the lifespan. Each plot includes information about the locus, the location and orientation of the genes in the region, the local estimates of recombination rate (in the right corner), and the LD estimates of surrounding SNPs with the index SNP (r^2^ values are estimated based on 1000 Genomes phase 3), which is indicated by colour (in the upper left corner).

One of the four loci identified in the single variant analysis reached genome-wide significance in the previous GWAS-MA on ADHD by Demontis and Walters et al.^6^, and all of them showed consistent direction of the effect between the two studies (Supplementary Table 8A). Significant loci reported by Demontis and Walters et al.^6^ showed nominal association with ADHD across the lifespan in the present study (Supplementary Table 8B and 8C), with single variant hits showing the same direction of the effect (Supplementary Table 8B).

Analyses conditioning on the index variant for the four ADHD-associated loci did not reveal new independent markers associated with ADHD. For each variant within the Bayesian credible sets obtained for these four loci, we searched for expression quantitative trait loci (eQTL) using data from blood^41^ and from a meta-analysis across different brain regions^42^. After integrating this information, credible sets for three of the four loci contained at least one eQTL within 1Mb of the index variant. The credible set on chromosome 6 included the index variant (rs183882582) and rs12197454. This variant, in LD with the index variant (r^2^ =0.56), was associated with the expression of *RSPH3* in blood and brain (P.adj<1.65E-05 and P.adj=2.36E-07, respectively) and with the expression of *VIL2* in blood (P.adj=3.21E-03). The credible set for the second most associated locus on chromosome 7 included 24 variants. The index variant, rs3958046, as well as additional variants in this credible set, were eQTLs for *CADPS2* in brain (maximum P.adj=2.91E-03). The credible set for the locus on chromosome 4 contained 50 variants, most of them located in or near *PCDH7*, but no eQTLs were identified. In the credible set for the locus on chromosome 3, which included 98 variants, the index variant, rs1920644, was associated with the expression of *KPNA4, IFT80*, and *KRT8P12* in brain (P.adj=1.16E-04, P.adj=1.40E-03, and P.adj=1.77E-03, respectively). Many other variants in this credible set were eQTLs for these genes and also for *TRIM59, OTOL1*, and/or *C3orf80* in brain (P.adj<0.05) (Supplementary Table 9 [in excel]).

In a summary-data-based Mendelian Randomization (SMR) analysis, we used summary data from the GWAS-MA of ADHD across the lifespan and the eQTL data in blood and brain from Westra et al.^41^ and Qi et al.^42^ to identify gene expression levels associated with ADHD. We found a significant association between ADHD across the lifespan and *RMI1* expression in blood after Bonferroni correction (P_SMR_=5.36E-06) (Supplementary Table 10 [in excel]). Results from the HEIDI test indicated that this finding was not an artifact due to linkage disequilibrium between eQTL and other ADHD associated variants (P_HEIDI_=0.60).

### Genetic correlation with other ADHD datasets and phenotypes

We found significant genetic correlations of ADHD in children and adults from the previous GWAS-MA^6^ (N=53,296) with persistent ADHD (rg=0.85, SE=0.04, P=5.49E-99), ADHD in childhood (rg=0.99, SE=0.03, P=5.02E-273), and ADHD across the lifespan (rg=0.98, SE=0.01, P<2.23E-308) (Supplementary Table 11). When we excluded sample overlap and considered the subset of new samples in our GWAS-MA on ADHD across the lifespan that were not included in the previous GWAS-MA by Demontis and Walters et al.^6^ (N=7,086), a significant genetic correlation was also obtained (rg=0.91, SE=0.35, P=8.70E-03).

We also observed significant genetic correlations between childhood ADHD symptom scores from a GWAS-MA in a population of children reported by the EAGLE consortium^43^ (N=17,666) and persistent ADHD (rg=0.65, SE=0.20, P=1.10E-03), ADHD in childhood (rg=0.98, SE=0.21, P=2.76E-06), and ADHD across the lifespan (rg=0.87, SE=0.19, P=4.80E-06). Similarly, significant genetic correlations between GWAS of self-reported ADHD status from 23andMe (N=952,652) and persistent ADHD (rg=0.75, SE=0.05, P=2.49E-45), ADHD in childhood (rg=0.63, SE=0.05, P=1.39E-42), and ADHD across the lifespan (rg=0.72, SE=0.04, P=4.86E-88) were observed (Supplementary Table 11).

We estimated the genetic correlation of persistent ADHD in adults, ADHD in childhood, and ADHD across the lifespan with all available phenotypes in LD-hub^44^, including ADHD-related traits and other psychiatric and neurological disorders. In total, results for 139 phenotypes passed the quality control parameters (h^2^ >0.1 and z-score>4) and 41 genetic correlations were significant after Bonferroni correction in both children and adults with persistent ADHD (Supplementary Table 12 [in excel]). The genetic correlations with ADHD were consistent across the lifespan, with similar patterns found in adulthood and childhood (Pearson’s r=0.89) (Figure 4A, Supplementary Table 12 [in excel]). The phenotypes showing the strongest genetic correlations with ADHD were traits related to academic performance, intelligence and risk-taking behaviors, including smoking and early pregnancy (Figure 4B).

**Figure 4.**
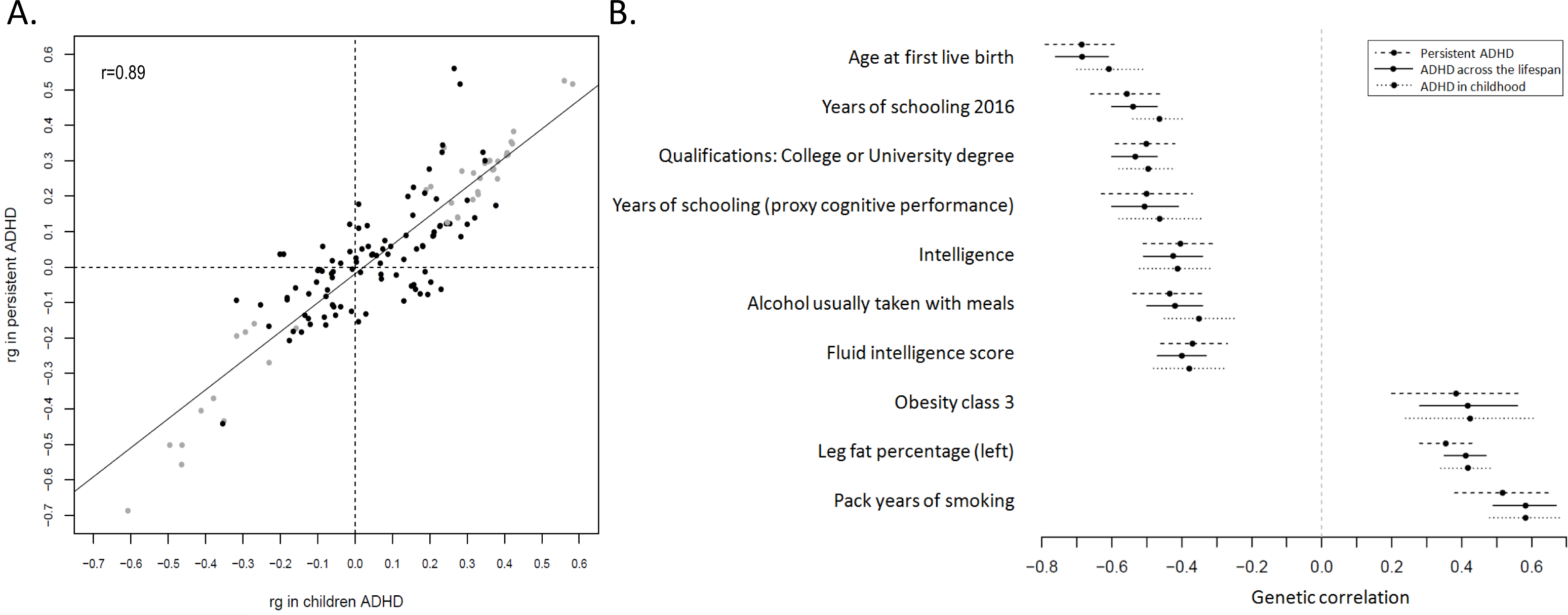
Genetic correlation of ADHD and several traits. (A) Black and grey dots represent genetic correlations (rg) for all traits considered (with h^2^ >0.1 and z-score>4) and for those traits which met Bonferroni correction in both children and adult ADHD groups, respectively. r indicates Pearson’s correlation coefficient. (B) The 10 strongest genetic correlations (with 95% confidence intervals) surpassing Bonferroni corrections in the children and persistent ADHD analysis are shown for each trait and ADHD.

## Discussion

In the current study, we set out to explore the contribution of common genetic variants to the risk of ADHD across the lifespan. Using the largest GWAS datasets available from the Psychiatric Genomics Consortium (PGC), the Lundbeck Foundation Initiative for Integrative Psychiatric Research (iPSYCH), and the International Multi-centre persistent ADHD CollaboraTion (IMpACT) consortia, we found evidence for a common genetic basis for ADHD in childhood and ADHD in adults that meet DSM-IV/ICD-10 criteria. We report a high genetic correlation between childhood and persistent ADHD in adulthood, and identified nine new loci associated with the disorder in single-variant and/or gene-based analyses.

We found a highly similar proportion of the heritability of ADHD explained by common variants in children (SNP-h^2^=0.21) and in adults (SNP-h^2^=0.20). This is consistent with the ADHD SNP-h^2^ estimate reported in the recent GWAS-MA (SNP-h^2^=0.22)^6^, which mainly (but not exclusively) included children with ADHD. Our finding is in line with multiple studies supporting the stability of ADHD’s heritability from childhood to adulthood^3-5^. The heritability results, together with the 0.81 genetic correlation found between children and adults with ADHD reinforce the hypothesis of the neurodevelopmental nature of persistent ADHD in adults. Consistently, the sign test and the PRS analysis confirmed the extensive overlap of common genetic risk variants for ADHD in childhood and adulthood. In addition, the result of the BUHMBOX analysis supported genetic similarities in ADHD across the lifespan with no evidence of a subset of patients enriched for persistent ADHD-associated alleles within the group of children.

Despite not having identified specific genetic contributions for ADHD in children or persistent ADHD, our results are not inconsistent with evidence suggesting changes in the genetic contribution to ADHD symptoms from childhood into adulthood, as described in previous twin studies in the general population^4,5,29,30^. Our study design and the still limited statistical power of the GWAS-MAs on ADHD may have facilitated the identification of the shared genetic basis rather than specific genetic factors for persistence. In addition, differences between such twin studies and the present study in the origin of the samples (population-based versus clinical) and/or discrepancies between self- and medical reports could also explain the reason for not identifying genetic variants associated specifically with childhood and/or persistent ADHD in our study. In addition, given that Chen et al.^45^ and Biederman et al.^46^ reported that persistence of ADHD into adulthood indexed stronger familial aggregation of ADHD, we cannot yet discard other types of genetic variation, such as rare mutations or copy number variation, playing a role in the different ADHD trajectories across the lifespan.

We also found strong and significant positive genetic correlations of ADHD ascertained in clinical populations of adults, children or both with other ADHD-related measures from general population samples, including the largest GWAS of self-reported ADHD status from 23andMe participants (N=952,652) and the GWAS-MA of childhood rating scales of ADHD symptoms in the general population^43^. In agreement with previous reports, these data suggest that a clinical diagnosis of ADHD in adults is an extreme expression of continuous heritable traits^6^ and that a single question about ever having received an ADHD diagnosis, as in the 23andMe sample, may be informative for molecular genetics studies.

Similar patterns of genetic correlation of ADHD with different somatic disorders and anthropometric, cognitive and educational traits were identified for children and adults with ADHD. These findings were highly similar to those observed in the recent GWAS-MA^6^ and further extend the existing hypothesis of a shared genetic architecture underlying ADHD and these traits to a lifespan perspective.

We report 13 loci in gene- and SNP-based analyses for childhood ADHD, adult ADHD and/or ADHD across the lifespan. Four ADHD-associated loci were previously identified by Demontis and Walters et al.^6^, which was expected due to the sample overlap between the two datasets (42,609 individuals shared out of the 49,560 in our study). The new loci identified in the present study mainly included genes involved in brain formation and function, such as *FEZF1*, a candidate for autism spectrum disorder implicated in the formation of the diencephalon^47,48^, *RSPH3*, which participates in neuronal migration in embryonic brain^49^, *CADPS2*, which has been associated with many psychiatric conditions due to its role in monoamine and neurotrophin neurotransmission^50-53^, *AMN*, which is involved in the uptake of vitamin B12^54,55^, essential for brain development, neural myelination, and cognitive function^56^, and *FBXL17*, which has previously been related to intelligence^57^.

In summary, the present cross-sectional analyses identify new genetic loci associated with ADHD and, more importantly, confirm that persistent ADHD in adults is a neurodevelopmental disorder that shows a high and significant genetic overlap with ADHD in children.

## Online methods

### Sample Description

A total of 19 GWAS of ADHD comprising 49,560 individuals (17,149 cases and 32,411 controls), provided by the Psychiatric Genomics Consortium (PGC), the Lundbeck Foundation Initiative for Integrative Psychiatric Research (iPSYCH), and the International Multi-centre persistent ADHD CollaboraTion (IMpACT), were analyzed in the present study. All participants were of European ancestry and provided informed consent and all sites had documented permission from local ethics committees. The meta-analysis on persistent ADHD included 22,406 individuals (6,532 ADHD adult cases and 15,874 controls) from nine datasets. The sample consisted of six datasets from the IMpACT consortium, two datasets from the PGC and the subset of adults from the iPSYCH cohort included in Demontis and Walters et al^6^. The meta-analysis on ADHD in childhood included 27,154 individuals (10,617 cases and 16,537 controls) from 10 datasets. The sample consisted of two new datasets from Brazil and Spain, seven datasets from the PGC and the subset of children from the iPSYCH cohort included in Demontis and Walters et al^6^. A total of 9,187 samples (4,281 cases and 4,906 controls) included in the iPSYCH cohort in Demontis and Walters et al.^6^ were not included in our GWAS-MA due to the distribution of children and adults across the different genotyping waves. Detailed information on each dataset is provided in Supplementary Table 1 and in the Supplementary Material.

### Genotyping, imputation, and quality control

Genotyping platforms and quality control (QC) filters for each of the 19 ADHD datasets are shown in Supplementary Table 1. Pre-imputation QC at individual and SNP level, principal component analyses for ancestry genetic outlier detection, and identification of related and/or duplicated individuals and gender discrepancies were performed using the Rapid Imputation and COmputational PIpeLIne (Ricopili) with the default settings (https://sites.google.com/a/broadinstitute.org/ricopili/). Non-European ancestry samples, related and duplicated individuals, and subjects with sex discrepancies were excluded. Phasing of genotype data was performed using SHAPEIT2 algorithm, and imputation for unrelated samples and trios was performed with MaCH, IMPUTE2, or MINIMAC3 (http://genome.sph.umich.edu/wiki/Minimac3) depending on software availability at the time of imputation^58-60^ (Supplementary Table 1). The European ancestry panels of the 1000 Genomes Project Phase 1 version 3 (v3) (April 2012) and Phase 3 version 5 (v5) (October 2014) using genome build hg19 were considered as references for the imputation of adult and children samples, respectively (ftp://ftp.1000genomes.ebi.ac.uk/vol1/ftp/).

### GWAS and meta-analyses

Imputed dosages and logistic regression analysis implemented in PLINK 1.9 were used assuming an additive model^61^. In the case-control studies, sex, the first 10 principal components, and other relevant covariates for each study were included (Supplementary Table 1). Summary statistics from each study were filtered prior to meta-analysis, excluding variants with minor allele frequency (MAF) <0.01 and imputation quality scores (INFO) ≤0.8. Inverse-variance weighted fixed-effects meta-analyses were conducted using METAL^62^, and results were filtered by effective sample size >70% of the total, defined as 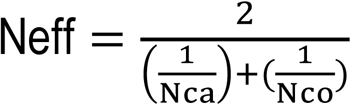^63^. After filtering, 7,366,995, 7,465,837 and 7,069,304 SNPs and insertion-deletion variants were included in the GWAS meta-analyses (GWAS-MA) of persistent ADHD, ADHD in childhood, and ADHD across the lifespan, respectively. Genomic inflation factors (λ and λ1000) and the ratio from LD score regression (the proportion of the inflation in the mean chi^2^ that the LD Score regression intercept ascribes to causes other than polygenic heritability) were calculated to check population stratification and technical biases. The genome-wide significance threshold was set at P<5.00E-08 to correct for multiple testing. Independent loci for variants exceeding the genome-wide significance threshold were defined based on clumping using PLINK 1.9. Variants that were ±250 kb away from the index variant (variant with smallest P-value in the region), with P-value<0.001 and with an estimated linkage disequilibrium (LD) of r^2^ >0.2 with the index variant were assigned to a clump (p_1_ =5.00E-08, p_2_ =0.001, r^2^ =0.2, kb=250). Manhattan and Forest plots were generated using the ‘qqman’ and ‘forestplot’ R packages (3.4.4 version of R), respectively. The LocusZoom software^64^ was used to generate regional association plots considering variants located ±500 kb from each index variant. r^2^ values between index and secondary variants were estimated based on the European ancestry 1000 Genomes Project Phase 3 reference panel.

### Conditional analysis

Conditional analyses for top-signals identified in the GWAS-MA of ADHD across the lifespan (±1,000 kb from each index variant position) were performed using the Genome-wide Complex Trait Analysis (GCTA) software^65^ and an in-house cohort of 3,727 individuals of European ancestry, imputed to the 1000 Genomes Project Phase 3 v5 reference panel (October 2014) as a reference for LD calculations.

### Bayesian credible set analysis

Credible sets of genetic variants that were 99% likely, based on posterior probability, to contain the causal variant, were defined using the method described by Maller et al.^66^ and implemented using a freely available R script (https://github.com/hailianghuang/FM-summary). We included variants located ± 250 kb away from the index variant, with P<1.00E-03 and with r2>0.2. The European ancestry 1000 Genomes Project Phase 3 v5 panel (October 2014) was used as reference for LD calculations.

### Functional characterization of genome-wide significant loci

Variant annotation integrator from UCSC (https://genome.ucsc.edu/cgi-bin/hgVai) was used to obtain functional annotation based on the Human hg19 genome build (Feb 2009). Credible causal sets of variants defined above were merged with the summary statistics of expression quantitative trait loci (eQTL) data from peripheral blood^41^ and with eQTL data from a meta-analysis across different brain regions (Brain-eMeta data from Qi et al.^42^). In the blood dataset, we used the false discovery rate (FDR) information provided^41^, and in the brain dataset we used the ‘stats’ R package to estimate the FDR (Benjamini & Hochberg) for variants included in any of the four credible sets. The statistical significance for both datasets was set using a threshold of 1.00E-03 for adjusted P-values. Summary data-based Mendelian Randomization (SMR)^67^ was used to test for association between gene expression levels and ADHD using summary statistics from the GWAS-MA of ADHD across lifespan and eQTL summary data from blood and brain datasets^41,42^. The heterogeneity in dependent instruments test (HEIDI) was used to assess whether the SMR findings were due to pleiotropy or linkage^67^.

### SNP-based heritability

The SNP-based heritability (SNP-h^2^) was estimated by single-trait LD score regression using summary statistics, HapMap 3 LD-scores, and considering default SNP QC filters (INFO>0.9 and MAF>0.01)^68^. Data of 1,113,287, 1,072,558, and 1,092,418 SNPs from the GWAS-MA of persistent ADHD, ADHD on childhood, and ADHD across the lifespan, respectively, were considered for the calculation of SNP-h^2^ estimates. Partitioning and enrichment of the heritability by functional categories was analyzed using the 24 main annotations (no window around the functional categories) described by Finucane et al^69^. Statistical significance was set using Bonferroni correction (P<2.08E-03).

### Gene-based and gene-set analyses

MAGMA software was undertaken for gene-based and gene-set association testing using summary data from our GWAS-MAs^70^. Variants were mapped to a gene if they were within 20 kb upstream or downstream from the gene according to dbSNP build 135 and NCBI 37.3 gene definitions. Genes in the MHC region (hg19:chr6:25-35M) were excluded from the analyses. LD patterns were estimated using the European ancestry 1000 Genomes Project Phase 3 v5 (October 2014) reference panel. Gene sets denoting canonical pathways were downloaded from MSigDB (http://www.broadinstitute.org/gsea/msigdb), which integrates Kyoto Encyclopedia of Genes and Genomes (KEGG) (http://www.genome.jp/kegg/), BioCarta (http://www.biocarta.com/), Reactome (https://reactome.org/) and Gene Ontology (GO) (http://www.geneontology.org/) resources. Bonferroni correction (P<2.77E-06 for 18,038 genes in persistent ADHD; P<2.75E-06 for 18,218 genes in childhood ADHD; P<2.79E-06 for 17,948 genes in ADHD across the lifespan) and 10,000 permutations were used for multiple testing correction in the gene-based and gene-set analyses, respectively.

### BUHMBOX analysis

The Breaking Up Heterogeneous Mixture Based On cross(X)-locus correlations (BUHMBOX) analysis^71^ was used to test whether the genetic correlation between persistent ADHD and ADHD in childhood was driven by subgroup heterogeneity, found when there is a subset of children enriched for persistent ADHD-associated alleles. Subgroup heterogeneity was tested in each childhood dataset considering two different SNP sets from the GWAS-MA of persistent ADHD, with P-value thresholds of P<5.00E-05 (62 LD-independent SNPs) and P<1.00E-03 (710 LD-independent SNPs). LD-independent variants with MAF>0.05 were defined by r^2^ >0.1 and a distance between them greater than 10,000 kb. BUHMBOX results were meta-analyzed using the standard weighted sum of z-score approach, where z-scores are weighted by the square root of the effective sample size 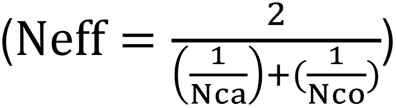^63^. The statistical power was calculated using 1,000 simulations, considering the ADHD children meta-analysis sample size (10,617 ADHD cases and 16,537 controls), the odds ratios and risk allele frequencies from the GWAS-MA of persistent ADHD and assuming 65% of heterogeneity proportion (π). The analyses of the sets of 62 and 710 variants had 98.4% and 100% of statistical power, respectively.

### Sign test

The direction of the effect of variants associated with ADHD in childhood was tested in persistent ADHD and vice versa, using strict clumping (r^2^ =0.05, kb=500, p_2_ =0.5) and different P-value thresholds (1.00E-07, 5.00E-07, 1.00E-06, 5.00E-06, 1.00E-05, 5.00E-05, 1.00E-04, and 5.00E-04). The concordant direction of effect was evaluated using a one sample test of the proportion with Yates’ continuity correction against a null hypothesis of P=0.50 with the ‘stats’ R package.

### Polygenic risk scoring

Non-ambiguous strand, independent SNPs (p_1_ =1, p_2_ =1, r^2^ =0.1, kb=250) with Neff>70% were selected from the GWAS-MA of ADHD in childhood at different P-value thresholds (P<0.001, 0.05, 0.1, 0.2, 0.3, 0.4, 0.5, and 1) to construct polygenic risk scores (PRSs) that were tested for association with persistent ADHD in each of the nine datasets using PRSice-2 (https://choishingwan.github.io/PRSice/). Best guess genotypes from the persistent ADHD datasets were filtered by excluding variants with MAF<0.01, INFO≤0.8 and missing rate >0.02, and only SNPs present in the childhood ADHD GWAS-MA results and in all the persistent ADHD studies were included in the analysis (N=32,584). Each study used the same covariates as included in the GWAS. Results from the nine PRS analyses at each P-value threshold, as well as results for all quintiles, were combined using the inverse-variance weighted meta-analysis.

### Genetic correlation

Cross-trait LD score regression with unconstrained intercept was used to calculate genetic correlations (rg) between pairs of traits, considering HapMap3 LD-scores, markers with INFO≥0.90, and excluding the MHC region (hg19:chr6:25-35M)^68^. Other ADHD datasets^6,43^ and phenotypes from the LD-hub centralized database^44^ (http://ldsc.broadinstitute.org) with heritability z-scores (observed heritability/observed standard error) >4 and with an observed heritability >0.1 were considered (N=139 out of 689 available traits). Statistical significance was set using Bonferroni correction (P<3.60E-04). Pearson’s correlation coefficient (Pearson’s r) was calculated between the genetic correlations of persistent ADHD with the phenotypes from the LD-hub and the genetic correlations of ADHD in childhood with the phenotypes from the LD-hub.

## Supporting information

Rovira_Supplementary_Material

Rovira_Supplementary_Table3

Rovira_Supplementary_Table8

Rovira_Supplementary_Table9

Rovira_Supplementary_Table11

Rovira_Supplementary_Table12

## Acknowledgements

P.R. is a recipient of a pre-doctoral fellowship from the Agència de Gestió d’Ajuts Universitaris i de Recerca (AGAUR), Generalitat de Catalunya, Spain (2016FI_B 00899). The iPSYCH project is funded by the Lundbeck Foundation (grant no R102-A9118 and R155-2014-1724) and the universities and university hospitals of Aarhus and Copenhagen. A.D.B. is also supported by the EU’s Horizon 2020 programme (grant no 667302, CoCA). Data handling and analysis was supported by NIMH (1U01MH109514-01 to Michael O’Donovan and Anders D. Børglum). High-performance computer capacity for handling and statistical analysis of iPSYCH data on the GenomeDK HPC facility was provided by the Centre for Integrative Sequencing, iSEQ, Aarhus University, Denmark (grant to Anders D. Børglum) and Center for Genomics and Personalized Medicine, Aarhus, Denmark. Over the course of this investigation, C.S.M. is a recipient of a Sara Borrell contract from the Instituto de Salud Carlos III, Ministerio de Economía, Industria y Competitividad, Spain (CD15/00199). N.R.M’s work was supported by the European Community’s Horizon 2020 Programme (H2020/2014 – 2020) under grant agreement n° 667302 (CoCA). K.P.L. and his team are supported by the Deutsche Forschungsgemeinschaft (DFG: CRU 125, CRC TRR 58 A1/A5, No. 44541416), the European Union’s Seventh Framework Programme under Grant No. 602805 (Aggressotype), the Horizon 2020 Research and Innovation Programme under Grant No. 728018 (Eat2beNICE) and 643051 (MiND), ERA-Net NEURON/RESPOND, No. 01EW1602B, ERA-Net NEURON/DECODE, No. 01EW1902 and 5-100 Russian Academic Excellence Project. In addition, this work was supported by the European College of Neuropsychopharmacology (ECNP Network “ADHD across the lifespan”). J.H. thanks Stiftelsen K.G. Jebsen, University of Bergen, the Western Norwegian Health Authorities (Helse Vest.) B.C. received financial support from the Spanish “Ministerio de Economía y Competitividad” (SAF2015-68341-R) and AGAUR (2017SGR738). The research leading to these results has also received funding from the European Union H2020 Program [H2020/2014-2020] under grant agreement n° 667302 (CoCA). J.A.R.Q. thanks patients and their families. M.S.A. is a recipient of a contract from the Biomedical Network Research Center on Mental Health (CIBERSAM), Madrid, Spain. M.R. is a recipient of a Miguel de Servet contract from the Instituto de Salud Carlos III, Spain (CP09/00119 and CPII15/00023). H.L. thanks to the Swedish research council. M.K.’s work is supported by the Dutch National Science Agenda NeurolabNL project (grant 400-17-602). E.Sprooten’s work is supported by a personal Hypatia grant from the Radboud University Medical Center. M.Hoogman received a Veni grant from of the Netherlands Organization for Scientific Research (NWO, grant number 91619115). The NeuroIMAGE study was supported by NIH Grant R01MH62873 (to Stephen V. Faraone), NWO Large Investment Grant 1750102007010 (to Jan Buitelaar), ZonMW grant 60-60600-97-193, NWO grants 056-13-015 and 433-09-242, and matching grants from Radboud University Nijmegen Medical Center, University Medical Center Groningen and Accare, and Vrije Universiteit Amsterdam. The research leading to these results also received support from the European Community’s Seventh Framework Programme (FP7/2007-2013) under grant agreement number 278948 (TACTICS) and under grant agreement number 602805 (Aggressotype).

This study is part of the International Multicentre persistent ADHD Collaboration (IMpACT; www.impactadhdgenomics.com). IMpACT unites major research centres working on the genetics of ADHD persistence across the lifespan and has participants in The Netherlands, Germany, Spain, Norway, the United Kingdom, the United States, Brazil and Sweden. Principal investigators of IMpACT are: Barbara Franke (chair), Andreas Reif (co-chair), Stephen V. Faraone, Jan Haavik, Bru Cormand, Josep Antoni Ramos-Quiroga, Philip Asherson, Klaus-Peter Lesch, Jonna Kuntsi, Claiton H.D. Bau, Jan K. Buitelaar, Stefan Johansson, Henrik Larsson, Alysa Doyle, and Eugenio H. Grevet. Their work is supported by the European Community’s Seventh Framework Programme (FP7/2007 – 2013) under grant agreement n° 602805 (Aggressotype) as well as from the European Community’s Horizon 2020 Programme (H2020/2014 – 2020) under grant agreements n° 643051 (MiND), n° 667302 (CoCA), and n° 728018 (Eat2beNICE). The work was also supported by the ECNP Network ‘ADHD across the Lifespan’ (https://www.ecnp.eu/research-innovation/ECNP-networks/List-ECNP-Networks/ADHD.aspx). B.F. received additional funding from a personal Vici grant of the Dutch Organization for Scientific Research (NWO; grant 016-130-669) and from a grant for the Dutch National Science Agenda (NWA) for the NeurolabNL project (grant 400 17 602). This paper represents independent research part funded by the National Institute for Health Research (NIHR) Biomedical Research Centre at South London and Maudsley NHS Foundation Trust and King’s College London. The views expressed are those of the authors and not necessarily those of any of the funding agencies, the NHS, the NIHR or the Department of Health and Social Care. S.V.F.’s work is supported by the European Union’s Seventh Framework Programme for research, technological development and demonstration under grant agreement no 602805, the European Union’s Horizon 2020 research and innovation programme under grant agreements No 667302 & 728018 and NIMH grants 5R01MH101519 and U01 MH109536-01. P.A.’s research is supported by the National Institute for Health Research (NIHR) Biomedical Research Centre for Mental Health, NIHR/MRC (14/23/17) and an NIHR Senior Investigator award (NF-SI-0616-10040). A.D. thanks the Stanley Center for Psychiatric Research. We thank the employees and research participants of 23andMe who made this study possible.

This study uses data from the ADHD Working Group of the Psychiatric Genomics Consortium (PGC), many of which contributed to this study and are named under the consortium author for this working group; additional members are at the time of submission of this manuscript: Anke R. Hammerschlag, Alexandra Philipsen, Alexandre Todorov, Alice Charach, Allison Ashley-Koch, Amaia Hervás, Ana Miranda, André Scherag, Anita Thapar, Anna Rommel, Anne Wheeler, Christine Cornforth, Danielle Posthuma, Elizabeth Corfield, Felecia Cerrato, Fernando Mulas, Franziska Degenhardt, Gláucia Chiyoko Akutagava Martins, Gun Peggy Strømstad Knudsen, Hans-Christoph Steinhausen, Herbert Roeyers, Hyo-Won Kim, Joanna Martin, Joel Gelernter, Joseph Sergeant, Juanita Gamble, Julia Pinsonneault, Jurgen Deckert, Kate Langley, Li Yang, Lindsey Kent, Manuel Mattheisen, Maria Jesús Arranz Calderón, Martin Steen Tesli, Meg Mariano, Michael Gill, Michael O’Donovan, Monica Bayes, Nick Martin, Niels Peter Ole Mors, Nigel Williams, Pak Sham, Patrick Sullivan, Patrick WL Leung, Paul Arnold, Paul Lichtenstein, Peter Holmans, Preben Bo Mortensen, Rachel Guerra, Raymond Walters, Richard Anney, Richard Ebstein, Ridha Joober, Sarah Anthony, Sarah Medland, Sarojini Sengupta, Søren Dinesen, Steve Nelson, Susan Smalley, Susann Scherag, Tammy Biondi, Tim Silk, Tinca Polderman, Tony Altar, Yanli Zhang-James, Yufeng Wang.

## Author contributions

### Conception or design of the work

P.R., D.D., C.S.M, T.Z., C.P.J., O.R., E.Sprooten, A.A.V., P.Asherson, B.C., S.V.F, J.H., S.E.J., K.P.L., M.Casas, A.D.B., B.F., J.A.R.Q., M.S.A., M.R.

### Acquisition, analysis or interpretation of the data

P.R., D.D., C.S.M, T.Z., M.K., N.R.M, H.W., I.G.M., M.P., L.V., L.A., V.R., M.Corrales, C.F., R.B., G.E.M., P.Almos, A.E.D., O.G., M.Hoogman, C.P.J, S.K.S., O.R., E.Svirin, T.S., ADHD Working Group of the Psychiatric Genomics Consortium, 23andMe Research Team, E.J.S.S.-B., P.Asherson, C.H.D.B., J.K.B., B.C., J.H., S.E.J., H.L., K.P.L., A.R., L.A.R., M.Casas, A.D.B, B.F., J.A.R.Q., M.S.A., M.R.

### Drafted the work or substantively revised it

P.R., D.D., C.S.M, P.Almos, E.H.G., A.H., M.Hutz, P.M.K., A.J.L., O.R., D.L.R., A.S.O., B.S.D.S, E.Svirin, T.S., E.J.S.S-B, P.Asherson, C.H.D.B., B.C., S.V.F., J.K., H.L., A.R., L.A.R., M.Casas, A.D.B., B.F., J.A.R.Q., M.S.A., M.R.

## Competing interests

V.R. has served on the speakers for Eli Lilly, Rubio and Shire in the last 5 years. She has received travel awards from Eli Lilly and Co. and Shire for participating in psychiatric meetings. The ADHD Program has received unrestricted educational and research support from Eli Lilly and Co., Janssen-Cilag, Shire, Rovi, Psious and Laboratorios Rubió in the past two years. M.Corrales received travel awards for taking part in psychiatric meetings from Shire. C.F. received travel awards for taking part in psychiatric meetings from Shire and Lundbeck. G.E.M. received travel awards for taking part in psychiatric meetings from Shire. E.S.S.-B. received speaker fees, consultancy, research funding and conference support from Shire Pharma. Consultancy from Neurotech solutions, Copenhagen University and Berhanderling, Skolerne & KU Leuven. Book royalties from OUP and Jessica Kingsley. Financial support – Arrhus Univeristy and Ghent University for visiting Professorship. Editor-in-Chief JCPP – supported by a buy-out of time to University of Southampton and personal Honorarium. King’s College London received payments for work conducted by P.A.: consultancy for Shire, Eli-Lilly, Novartis and Lundbeck; educational and/or research awards from Shire, Eli-Lilly, Novartis, Vifor Pharma, GW Pharma, and QbTech; speaker at events sponsored by Shire, Eli-Lilly, Janssen-Cilag and Novartis. J.K.B., has been in the past 3 years a consultant to / member of advisory board of / and/or speaker for Shire, Roche, Medice, and Servier. He is not an employee of any of these companies, and not a stock shareholder of any of these companies. He has no other financial or material support, including expert testimony, patents, royalties. In the past year, Dr. S.V.F. received income, potential income, travel expenses continuing education support and/or research support from Tris, Otsuka, Arbor, Ironshore, Shire, Akili Interactive Labs, Enzymotec, Sunovion, Supernus and Genomind. With his institution, he has US patent US20130217707 A1 for the use of sodium-hydrogen exchange inhibitors in the treatment of ADHD. He also receives royalties from books published by Guilford Press: Straight Talk about Your Child’s Mental Health, Oxford University Press: Schizophrenia: The Facts and Elsevier: ADHD: Non-Pharmacologic Interventions. He is principal investigator of www.adhdinadults.com. J.K. has given talks at educational events sponsored by Medice; all funds are received by King’s College London and used for studies of ADHD. H.L. has served as a speaker for Evolan Pharma and Shire and has received research grants from Shire; all outside the submitted work. K.P.L. served as a speaker for Eli Lilly and received research support from Medice, and travel support from Shire, all outside the submitted work. L.A.R. reported receiving honoraria, serving on the speakers’ bureau/advisory board, and/or acting as a consultant for Eli-Lilly, Janssen-Cilag, Novartis, and Shire in the last 3 years; receiving authorship royalties from Oxford Press and ArtMed; and receiving travel awards from Shire for his participation in the 2015 WFADHD meetings and from Novartis to take part of the 2016 AACAP meeting. The ADHD and juvenile bipolar disorder outpatient programs chaired by him received unrestricted educational and research support from the following pharmaceutical companies in the last 3 years: Janssen-Cilag, Novartis, and Shire. sent by angelica: L.A.R. reported receiving honoraria, serving on the speakers’ bureau/advisory board, and/or acting as a consultant for Eli-Lilly, Janssen-Cilag, Novartis, and Shire in the last 3 years; receiving authorship royalties from Oxford Press and ArtMed; and receiving travel awards from Shire for his participation in the 2015 WFADHD meetings and from Novartis to take part of the 2016 AACAP meeting. The ADHD and juvenile bipolar disorder outpatient programs chaired by him received unrestricted educational and research support from the following pharmaceutical companies in the last 3 years: Janssen-Cilag, Novartis, and Shire. M.Casas has received travel grants and research support from Eli Lilly and Co., Janssen-Cilag, Shire and Lundbeck and served as consultant for Eli Lilly and Co., Janssen-Cilag, Shire and Lundbeck. B.F. has received educational speaking fees from Medice and Shire. J.A.R.Q. was on the speakers’ bureau and/or acted as consultant for Eli-Lilly, Janssen-Cilag, Novartis, Shire, Lundbeck, Almirall, Braingaze, Sincrolab, Medice and Rubió in the last 5 years. He also received travel awards (air tickets + hotel) for taking part in psychiatric meetings from Janssen-Cilag, Rubió, Shire, Medice and Eli-Lilly. The Department of Psychiatry chaired by him received unrestricted educational and research support from the following companies in the last 5 years: Eli-Lilly, Lundbeck, Janssen-Cilag, Actelion, Shire, Ferrer, Oryzon, Roche, Psious, and Rubió. Members of the 23andMe Research Team are current or former employees of 23andMe, Inc. and hold stock or stock options in 23andMe.

## Data availability

The full GWAS-MA summary statistics for the three meta-analyses will be available for download in the https://www.med.unc.edu/pgc/results-and-downloads webpage. The full GWAS summary statistics for the 23andMe ADHD data set will be made available through 23andMe to qualified researchers under an agreement with 23andMe that protects the privacy of the 23andMe participants. Please visit research.23andme.com/collaborate/#publication for more information and to apply to access the data.

