## Supplementary material for "Shared genetic background between children and adults with attention deficit/hyperactivity disorder": Rovira_Supplementary_Table12

*Study description*

**Persistent ADHD**

**Brazil**

This sample comprises 417 white Brazilians of European descent ascertained from the ADHD Outpatient Program in the adult division at Hospital de Clínicas de Porto Alegre (HCPA), Brazil. All individuals were 18 years or older and fulfilled ADHD diagnosis criteria both currently and during childhood (retrospectively). Diagnoses of ADHD followed DSM-IV criteria[1] and were carried out by trained psychiatrists through the application of a semi-structured interview, the Portuguese version of the Schedule for Affective Disorders and Schizophrenia for School-Age Children–Epidemiologic version (K-SADS-E), adapted to adults as described by Grevet et al[2]. Other lifetime psychiatric comorbidities were assessed by the Structured Clinical Interview for DSM-IV Axis I Disorders (SCID-I)[3]. Subjects presenting a clinically significant history of neurological disease (e.g., delirium, dementia, epilepsy, head trauma, multiple sclerosis), past or present symptoms of psychosis and/or an estimated IQ< 70 were excluded. The study sample also included 463 Brazilian adult controls ascertained at the blood donation center of HCPA. The inclusion criteria were (A) being Brazilian of European descent and (B) aged 18 years or older. The exclusion criteria were: (A) positive screening for ADHD in the 6-question adult self-report scale – V1.1 (ASRS-V1.1) screener[4], (B) evidence of a clinically significant neurological disease that might affect cognition (e.g., delirium, dementia, epilepsy, head trauma, and multiple sclerosis), and (C) current or past history of psychosis. Other psychiatric disorders were assessed by using the structured clinical interview for DSM-IV screening module – SCID-I for the Axis I psychiatric[5]. DNA was extracted from peripheral blood. The study was carried out in accordance with the Declaration of Helsinki, and all participants signed an informed consent form previously approved by the institutional review board of the hospital (No. 00000921).

**Denmark**

Biological material from the included individuals was obtained from the Danish Newborn Screening Biobank at Statens Serum Institute. This biobank contains blood spots (Guthrie cards) from all newborn babies that have been born in Denmark since 1981. This nationwide biobank can be linked with Danish registers through the unique personal identification number[6] (CPR-number), which is assigned to all live-born babies in Denmark and used in all contacts with the public sector, including the health care system. Persistent ADHD cases were identified in a nationwide population-based case-cohort sample selected from a baseline birth cohort comprising all singletons born in Denmark between May 1, 1981, and December 31, 2005, who were residents in Denmark on their first birthday and who have a known mother (N= 1,472,762). Cases were diagnosed by psychiatrists at in- or out-patient clinics, predominantly the latter according to the ICD10 criteria (F90.0 diagnosis code), identified using the Danish Psychiatric Central Research Register[7] (DPCRR). Diagnoses were given in 2016 or earlier for individuals at least 1 year old. Cases were considered as "persistent ADHD" cases if they had received the diagnosis of ADHD when they were older than 18 years. Controls were randomly selected from the same nationwide birth cohort and not diagnosed with ADHD (F90.0). Controls used in the GWAS of persistent ADHD were older than 18 years in 2016. DNA was extracted from dried blood spot samples and whole genome was amplified in triplicates as described previously[8,9]. Processing of DNA, genotyping and genotype calling as well as imputing of genotypes of the iPSYCH-ADHD sample were carried out as a part of the genotyping of the full iPSYCH sample, which in total consists of around 79,492 individuals. The data processing was done in 23 waves of approximately 3,500 individuals each. In order to control for potential batch effects, we included "wave" as a covariate. Following genotyping all data processing, quality control, and downstream analyses were performed at secured servers in Denmark at the GenomeDK high performance-computing cluster (http://genome.au.dk). The study was approved by the Danish Data Protection Agency and the Scientific Ethics Committee in Denmark.

**Germany**

Patients with a diagnosis of persistent ADHD were recruited by experienced psychiatrists at the University of Würzburg (Würzburg, Germany). Unrelated in– and outpatients of self-reported central-European descent completed a semi-structured clinical interview according to DSM-IV[3]. Inclusion criteria were onset before the age of 7 years, life-long persistence, current diagnosis and age of recruitment between 18 and 65 years. Patients with a lifetime diagnosis of bipolar disorder (BPD), schizophrenia, significant neurologic co-morbidity, mental retardation, substance-induced disorders (before detoxification/withdrawal), or other somatic disorders, suggesting organic psychosis or with an intelligence quotient (IQ) below 80 were excluded from analysis. For a more detailed sample description, please confer previous publications[10,11]. The controls consisted of 329 anonymous blood donors, not screened for psychiatric disorders, but free of medication and further 733 healthy volunteers, screened for psychiatric disorders. The study was approved by the Ethic Committee of the University of Würzburg (Würzburg, Germany). All participants thus were informed and gave written informed consent according to the principles of the Helsinki declaration. Genomic DNA was isolated from frozen venous EDTA-blood by a standard de-salting method.

**The Netherlands**

The ADHD patients and healthy subjects were recruited from the department of Psychiatry of the Radboud University Nijmegen Medical Centre and through advertisements. Patients were included if they met DSM-IV-TR criteria[5] for ADHD in childhood as well as adulthood. All subjects were assessed using the Diagnostic Interview for Adult ADHD (DIVA)[12]. In order to obtain information about ADHD symptoms and impairment in childhood, additional information was obtained from parents and school reports, whenever possible. The Structured Clinical Interview for DSM-IV Criteria (SCID-I)[3] was used for co-morbidity assessment. Assessments were carried out by trained professionals (psychiatrist or psychologists). Exclusion criteria for participants were psychosis, addiction in the last 6 months, current major depression (assessed with SCID-I)[3], full-scale IQ estimate<70 (Wechsler Adult Intelligence Scale-III), neurological disorders, sensorimotor handicaps, non-Caucasian ethnicity and medication use other than psychostimulants or atomoxetine. Additional exclusion criteria for healthy subjects were a current or past neurological or psychiatric disorder according to SCID-I[3]. This study was approved by the regional ethics committee. Written informed consent was obtained from all participants. Genomic DNA was extracted as specified by the manufacturer.

**Norway I and II**

The adult ADHD cases were recruited through a Norwegian national medical registry that was established to harmonize diagnostic assessment and treatment of adult ADHD across the nation. Between 1997 and 2005, all adult patients (older than 18 years) in Norway who were to receive treatment with central stimulants had to be evaluated and approved by one of three national expert committees for ADHD or hyperkinetic disorder. Clinicians first examined and diagnosed their patients according to nationally recommended guidelines. These guidelines included systematic assessment of ADHD diagnostic criteria, developmental history, physical examination, evaluation of comorbidity, and, where possible, information from collateral informants. The expert committees reviewed the data before a definitive diagnosis was established. The diagnosis of ADHD was made according to the International Classification of Diseases (ICD-10) research criteria, with two modifications: allowing the inattentive subtype in the Diagnostic and Statistical Manual of Mental Disorders as sufficient for the diagnosis and allowing for the presence of comorbid psychiatric disorders, as long as the diagnostic symptoms of ADHD were present before the appearance of the comorbid disorder. This diagnostic assessment strategy was chosen as a compromise between the fact that ICD-10 was the official diagnostic system in Norway and the request to have an assessment comparable with the DSM-IV standards[13]. The Norwegian patient sample also included adult patients participants diagnosed after 2005. These were also recruited by psychologists and psychiatrists at out-patient clinics, using DSM-IV criteria, according to national guidelines. A minority of the cases with persistent ADHD (8.3%) included in this study were also officially diagnosed with ADHD before 18 years of age. For this group, the average age at inclusion was 27.2 years, compared to 32.6 years for the ADHD cases first diagnosed as adults. The control group was recruited from the database of the Medical Birth Registry of Norway. This registry includes all people born in Norway after January 1, 1967. The geographical distribution was similar for patients and controls, with the majority coming from the Bergen and Oslo areas, but all regions of Norway were represented. All participants provided either blood or saliva samples for DNA extraction. All participants provided signed informed consent form. The study was approved by the Norwegian regional medical research ethics committee West (IRB #3 FWA00009490, IRB00001872). The recruitment protocols and clinical features of the samples have been described previously[13-15].

**Spain I, II and III**

All subjects were evaluated and recruited prospectively from a restricted geographic area in a specialized out-patient program for Adult ADHD and by a single clinical group at Hospital Universitari Vall d'Hebron of Barcelona (Spain). Patients were referred to the program from primary care centers and adult community mental health services. The study was approved by the Clinical Research Ethics Committee (CREC) of Hospital Universitari Vall d'Hebron, all methods were performed in accordance to the relevant guidelines and regulations and written informed consent was obtained from all subjects before inclusion into the study. The clinical assessment was conducted by structured interviews and self-reported questionnaires in two different steps: (i) Assessment of ADHD diagnosis based on symptomatology using the Conner's Adult ADHD Diagnostic Interview for DSM-IV (CAADID)[16] by a psychiatrist and, (ii) Assessment of the severity of ADHD symptoms, the levels of impairment and the presence of comorbid disorders by a psychologist to increase the diagnostic accuracy and reduce the likelihood of misdiagnosis with the Conners ADHD Rating Scale (CAARS)[17], the ADHD Rating Scale (ADHD-RS)[18], the Clinical Global Impression (CGI)[19], the Wender Utah Rating Scale (WURS)[20], the Sheehan Disability Inventory (SDS)[21], and the Structured Clinical Interview for DSM-IV Axis I and II Disorders (SCID-I and SCID-II)[3]. Afterwards, the psychiatrist and psychologist integrated the clinical information and self-reports for the valid assessment of symptoms and impairments. Given the clinical diagnosis of ADHD, in case of discordance between different raters of ADHD symptoms or inconsistencies between reporters in responses to items measuring similar symptoms, the clinician-identified symptoms on the CAADID prevailed. The diagnosis of ADHD was assessed with the CAADID structured interview, which is divided into Part I and Part II that were administered separately: (i) the CAADID Part I which is a Patient History administered as a clinical interview and (ii) the CAADID Part II which is a Diagnostic Criteria Interview used to assess ADHD according to DSM-IV criteria. Diagnosis of ADHD was confirmed when patients presented six or more symptoms of inattention and/or six or more symptoms of hyperactivity–impulsivity, which persisted for at least 6 months to a degree that is considered maladaptive and inconsistent with the developmental level. When possible, a collateral interview with a family member was performed during the clinical interview, which provides valuable information that the patient may not self-report. In addition to the clinical diagnosis, severity of ADHD symptoms and the level of impairment were assessed using self-report questionnaires that include: (i) the CAARS Scalelong version gathered from the patient (self-report; CAARS-S:L) and an observer (secondary informant; CAARS-O:L); the ADHD-RS and the CGI to measure the severity of ADHD symptoms in adulthood, (ii) the WURS to retrospectively assess ADHD symptoms in childhood and (iii) the SDS to assess the level of functional impairment. Finally, differential diagnosis and assessment of comorbidities were assessed with SCID-I and SCID-II[3]. Exclusion criteria was IQ<70; lifelong and current history of mood, psychotic, anxiety, substance abuse, and DSM-IV axis II disorders; pervasive developmental disorders; a history or the current presence of a condition or illness, including neurologic, metabolic, cardiac, liver, kidney, or respiratory disease; a chronic medication of any kind; birth weight ≤ 1.5 kg; and other neurological or systemic disorders that might explain ADHD symptoms. The control sample consisted of unrelated healthy blood donors matched by sex with the clinical group. Individuals with ADHD symptomatology were excluded under the following criteria: 1) having been diagnosed with ADHD previously and 2) answering positively to the life-time presence of the following ADHD symptoms: a) often has trouble in keeping attention on tasks, b) usually loses things needed for tasks, c) often fidgets with hands or feet or squirms in seat, and d) often gets up from seat when remaining in seat is expected. Genomic DNA of all participants was isolated from peripheral blood leukocytes by a salting out procedure.

### Childhood ADHD

#### Brazil

The Brazilian clinical sample included 259 ADHD children and adolescents and their biological parents recruited from the ADHD Outpatient Program (ProDAH) at the Hospital de Clínicas de Porto Alegre (HCPA). The diagnoses of ADHD and comorbidities were achieved through a three-step process: application of semistructured interviews (Schedule for Affective Disorders and Schizophrenia for School-

Age Children - Present and Lifetime Version, K-SADS-PL)[22], diagnostic discussion in a clinical committee, and clinical evaluation with the child and his/her parents. Experienced child psychiatrists confirmed all generated diagnoses. The Swanson, Nolan, and Pelham Scale-Version IV (SNAP-IV)[23] was also applied to patients by child psychiatrists. IQ was estimated through Wechsler Intelligence Scale for Children—Third Edition (WISC III) performed by trained psychologists[24,25]. The Ethics Committee of HCPA approved the study protocol. Parents provided written informed consent and children and adolescents provided verbal assent to participate.

**Denmark**

Biological material from the included individuals was obtained from the Danish Newborn Screening Biobank at Statens Serum Institute. This biobank contains blood spots (Guthrie cards) from all newborn babies that have been born in Denmark since 1981. This nationwide biobank can be linked with Danish registers through the unique personal identification number[6] (CPR-number), which is assigned to all live-born babies in Denmark and used in all contacts with the public sector, including the health care system. Persistent and childhood ADHD cases were identified in a nationwide population-based case-cohort sample selected from a baseline birth cohort comprising all singletons born in Denmark between May 1, 1981, and December 31, 2005, who were residents in Denmark on their first birthday and who have a known mother (N = 1,472,762). Cases were diagnosed by psychiatrists at in- or out-patient clinics, predominantly the latter according to the ICD10 criteria (F90.0 diagnosis code), identified using the Danish Psychiatric Central Research Register[7] (DPCRR). Diagnoses were given in 2016 or earlier for individuals at least 1 year old. Cases were considered as “childhood ADHD” if they received the diagnosis when they were less than 18 years old. Controls were randomly selected from the same nationwide birth cohort and not diagnosed with ADHD (F90.0). Controls used in the GWAS of childhood ADHD were less than 18 years old in 2016. DNA was extracted from dried blood spot samples and whole genome amplified in triplicates as described previously[8,9]. Processing of DNA, genotyping and genotype calling as well as imputing of genotypes of the iPSYCH-ADHD sample were carried out as a part of the genotyping of the full iPSYCH sample, which in total consists of around 79,492 individuals. The data processing was done in 23 waves of approximately 3,500 individuals each. In order to control for potential batch effects, we included “wave” as a covariate in the GWAS. Following genotyping all data processing, quality control, and downstream analyses were performed at secured servers in Denmark at the GenomeDK high performance-computing cluster (http://genome.au.dk). The study was approved by the Danish Data Protection Agency and the Scientific Ethics Committee in Denmark.

**Spain**

The study was approved by the Clinical Research Ethics Committee (CREC) of Hospital Universitari Vall d'Hebron, all methods were performed in accordance to the relevant guidelines and regulations and written informed consent was obtained from parents/caregivers before inclusion into the study. Participants were clinically interviewed using clinical history and the Diagnostic Interview Schedule for Affective Disorders and Schizophrenia for School-Age Children 6-18 years–Lifetime Version DSM-IV (K-SADS-PL)[22]. Subjects were required to satisfy full Diagnostic and Statistical Manual of Mental Disorders (DSM)-IV criteria for ADHD, be <18 years of age, Spanish of Caucasian origin and have never been treated for ADHD. Clinical assessment Diagnosis of ADHD and comorbidities were established by child psychiatrists blind to patients’ genotypes. General psychiatric symptomatology was evaluated based on the Achenbach System of Empirically Based Assessment (ASEBA)[26], the parent-reported Child Behavior Checklist (CBCL)[27], and the Teacher Report Form (TRF)[28]. Furthermore, families and teachers were interviewed with the parent and teacher versions of the Strengths and Difficulties Questionnaire (SDQ)[29], respectively. After the assessment, 430 subjects met diagnostic criteria for ADHD. Parents completed a general questionnaire on socio-demographic information and clinical data regarding their child and family psychiatric history. Intellectual level (IQ) was assessed with the Wechsler Intelligence Scale for children-IV (WISC-IV)[30]. Patients with an IQ<70 or having pervasive developmental disorders were not eligible for this study. Additional exclusion criteria included schizophrenia or other psychotic disorders; adoption; sexual or physical abuse; birth weight<1.5 kg; any significant neurological or systemic disease that might explain ADHD symptoms. Comorbid oppositional defiant disorder, conduct disorder, depression and anxiety disorders were allowed unless determined to be the primary cause of ADHD symptomatology. The control sample consisted of 639 unrelated healthy blood donors matched by sex with the clinical group. Individuals with ADHD symptomatology were excluded under the following criteria: 1) having been diagnosed with ADHD previously and 2) answering positively to the life-time presence of the following ADHD symptoms: a) often has trouble in keeping attention on tasks, b) usually loses things needed for tasks, c) often fidgets with hands or feet or squirms in seat, and d) often gets up from seat when remaining in seat is expected. Genomic DNA of all participants was isolated from peripheral blood leukocytes by a salting out procedure.

**Cohorts from the Psychiatric Genomics Consortium (PGC):**

Extensive information of the samples included in the CHOP (USA), IMAGE-I (Europe), PUWMa (USA), Canada (Toronto), UK (Cardiff), Germany, IMAGE-II (Europe and USA) cohorts used in the present study is available elsewhere[31].

**<u>Other ADHD datasets</u>**

**23andMe, self-reported ADHD diagnoses**

The 23andMe sample consists of research participants of the personal genetics company 23andMe, Inc., who submitted saliva samples and consented to participate in research. Participants answered questions online about their ADHD history according to 23andMe's human subjects protocol, which was reviewed and approved by Ethical & Independent Review Services, an AAHRPP-accredited institutional review board. The ADHD phenotype was defined by using the following survey questions: 1) Have you ever been diagnosed by a doctor with any of the following psychiatric conditions? (options for "Attention deficit disorder (ADD) or Attention deficit hyperactivity disorder (ADHD): Yes, No, I don't know); 2) Have you ever been diagnosed with attention-deficit disorder without hyperactivity (ADD)? (answers: Yes, No, I'm not sure); 3) Have you ever been diagnosed or treated for any of the following conditions? (options for "Attention deficit disorder (ADD)": Yes, No, I'm not sure); 4) Have you ever been diagnosed or treated for any of the following conditions? (options for "A mental health or psychiatric condition": Yes, No, I'm not sure); 5) What mental health problems have you had? Please check all that apply (checkbox: "Attention deficit hyperactivity disorder (ADHD)" or "Attention deficit disorder (ADD)"); 6) Have you ever been diagnosed with or treated for attention deficit disorders (ADHD or ADD)? (answers: Yes, No, I'm not sure); 7) Have you ever been diagnosed with or treated for any of the following conditions? Anxiety, Attention deficit disorders, Bipolar disorder/maniac depression, Depression, Eating disorder (such as anorexia or bulimia) (answers: Yes, No, I'm not sure); 8) In the last 2 years, have you been newly diagnosed with or started treatment for any of the following conditions? (options for "Attention deficit disorder (ADD)": Yes, No, I'm not sure). Research participants of European ancestry with successful genotypes genome-wide were used in the study. Genotyping was performed on the Illumina HumanHap550+ BeadChip (1,476 cases, 14,469 controls), Illumina OmniExpress+ BeadChip (16,853 cases, 147,299 controls), and a full custom array (78,008 cases, 694,547 controls). Imputation was performed against a combined reference panel consisting of 1000 Genomes Project Phase 3 and UK10K data. GWAS for the ADHD phenotype were performed with age, sex, the first five principal components, and genotyping platform as covariates. The 23andMe summary statistics were verified to be consistent with genome build hg19.

**EAGLE, ADHD symptom scores**

The EArly Genetics and Lifecourse Epidemiology (EAGLE) consortium includes population-based birth cohorts from Europe, Australia, and the United States (http://www.wikigenes.org/e/art/e/348.html). The consortium focuses on a wide range of phenotypes in childhood including traits related to cognition and behavior e.g. aggression[32], astma allergy and atopy[33] and postnatal growth[34]. In the study of ADHD symptoms, nine EAGLE cohorts where included with available ADHD symptom scores in childhood (age at measurement <13 years). An overview of the nine cohorts included in the EAGLE meta-analysis is provided in Middeldorp et al[35]. In order to assess ADHD symptoms different instruments were used across cohorts, including the Attention Problems scale of the Child Behavior Checklist (CBCL)[27] and the Teacher Report Form (TRF)[28], the Hyperactivity scale of the Strengths and Difficulties Questionnaire (SDQ)[29], and the DSM-IV ADHD items as, for example, included in the Conners Rating Scale[17]. For the meta-analysis, one phenotype was selected from each cohort. Based on the phenotype that was most available, school age ratings were chosen over preschool-age ratings, parent ratings over teacher ratings, and the measurement instrument with the largest information density was preferred over the other instruments[35].

For the present study, we used the summary data of the meta-analysis of ADHD symptoms available for downloading in https://www.med.unc.edu/pgc/results-and-downloads.

**Supplementary Figure 1. Q-Q plots for GWAS meta-analysis of (A) persistent ADHD, (B) ADHD in childhood and (C) ADHD across the lifespan.**

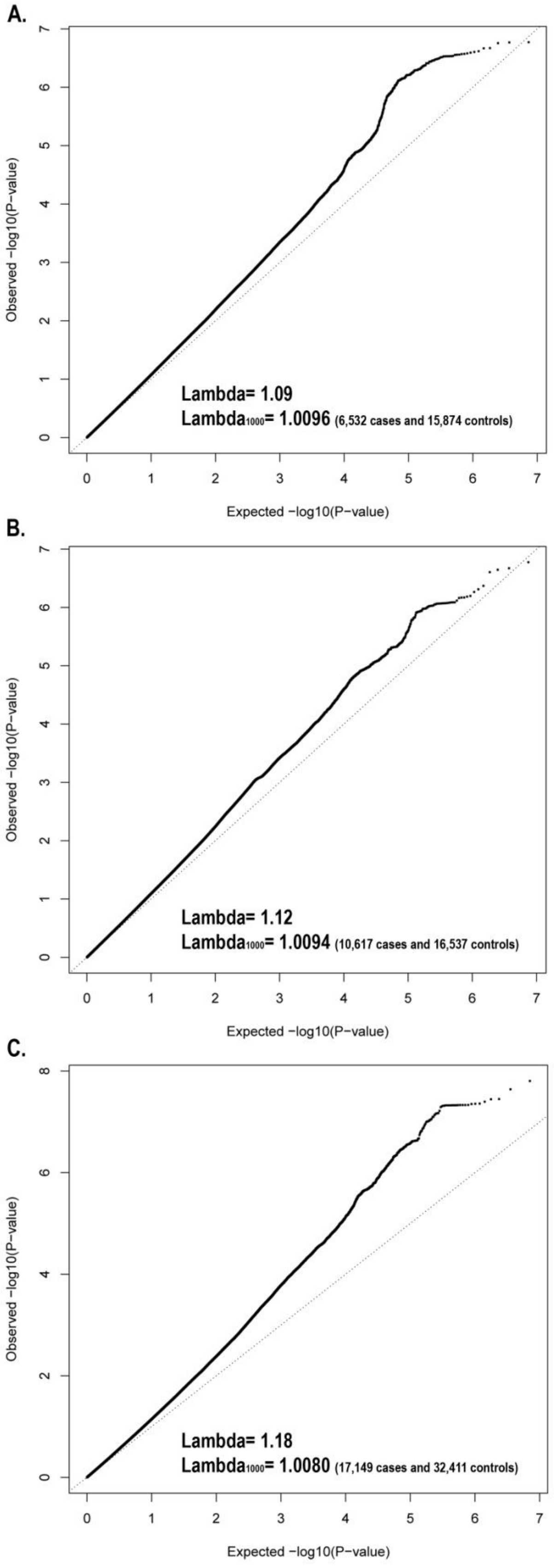

**Supplementary Figure 2. Partitioned heritability by functional annotations for (A) persistent ADHD, (B) ADHD in childhood and (C) ADHD across the lifespan.** Enrichment of SNP heritability in 24 functional annotations defined by Finucane et al [36]. Error bars represent 95% confidence intervals. Nominal significant enriched categories are labeled with an asterisk (*) and darker grey bars indicate significant enrichment after Bonferroni correction (P<2.08E-03).

**A.**

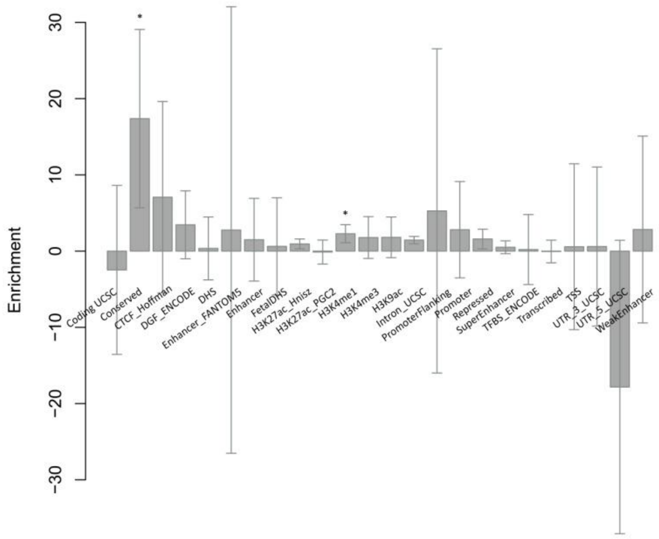


**B.**

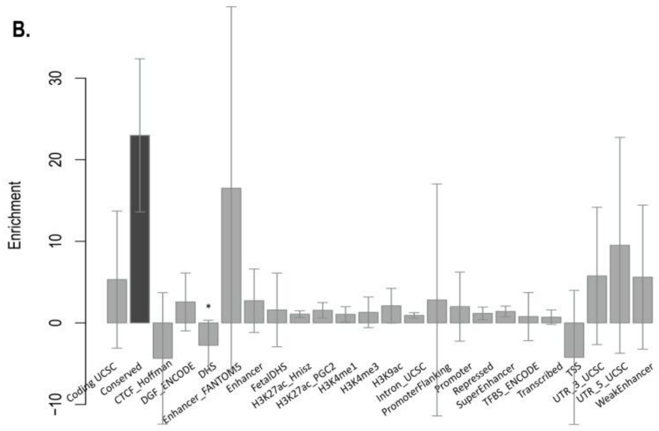


**C.**

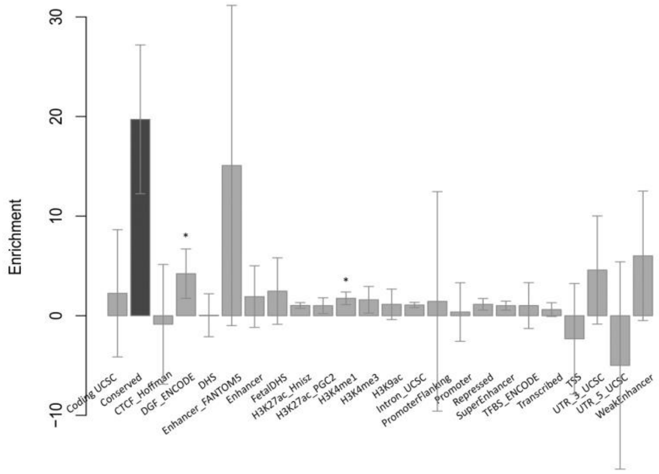

**Supplementary Figure 3. Forest plots for the genome-wide significant loci identified in the GWAS meta-analysis of ADHD across the lifespan.** Each plot (A-D) provides a visualization of the effect size estimates for each study and for the summary meta-analysis. The 95% confidence intervals are included for the estimates.

**A.**

**rs183882582**

| Study | Cases | Controls | OR |
|---|---|---|---|
| Germany adults | 552 | 1062 | 1.27 |
| Norway I | 305 | 208 | 1.89 |
| Norway II | 390 | 354 | 2.25 |
| Denmark adults | 3841 | 11707 | 1.42 |
| PUWMa, USA | 563 | 563 | 2.33 |
| Germany children | 487 | 1290 | 1.29 |
| IMAGE−I | 700 | 700 | 1.39 |
| IMAGE−II | 624 | 1755 | 2.04 |
| CHOP, USA | 262 | 262 | 1.13 |
| UK | 721 | 5081 | 1.39 |
| Brazil children | 259 | 259 | 0.88 |
| Denmark children | 6462 | 5879 | 1.35 |
| **Summary** | **15166** | **29120** | **1.42** |

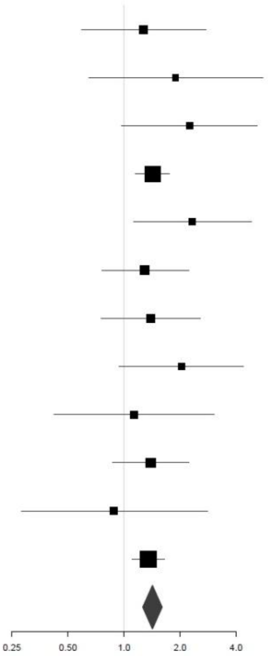


**B.**

**rs3958046**

| Study | Cases | Controls | OR |
|---|---|---|---|
| Germany adults | 552 | 1062 | 1.17 |
| Norway I | 305 | 208 | 0.95 |
| Norway II | 390 | 354 | 1.23 |
| Brazil adults | 417 | 463 | 1 |
| Spain I | 418 | 428 | 1 |
| Spain II | 126 | 126 | 1.77 |
| Spain III | 359 | 1414 | 1.3 |
| Denmark adults | 3841 | 11707 | 1.07 |
| PUWMa, USA | 563 | 563 | 1.05 |
| Canada | 109 | 109 | 0.88 |
| Germany children | 487 | 1290 | 1.15 |
| IMAGE−I | 700 | 700 | 1.09 |
| IMAGE−II | 624 | 1755 | 1.15 |
| CHOP, USA | 262 | 262 | 1.27 |
| UK | 721 | 5081 | 1.22 |
| Brazil children | 259 | 259 | 1.09 |
| Spain children | 430 | 639 | 1.03 |
| Denmark children | 6462 | 5879 | 1.05 |
| **Summary** | **17025** | **32299** | **1.09** |

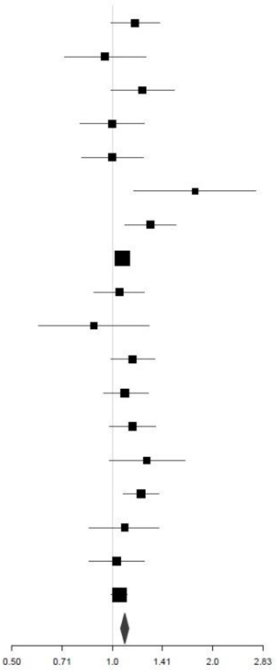

C.

**rs200721207**

| Study | Cases | Controls | OR |
|---|---|---|---|
| Germany adults | 552 | 1062 | 1.2 |
| The Netherlands | 124 | 112 | 1 |
| Norway I | 305 | 208 | 0.79 |
| Norway II | 390 | 354 | 1.05 |
| Brazil adults | 417 | 463 | 1.14 |
| Spain I | 418 | 428 | 1.08 |
| Spain II | 126 | 126 | 0.73 |
| Spain III | 359 | 1414 | 1.11 |
| Denmark adults | 3841 | 11707 | 1.09 |
| Germany children | 487 | 1290 | 1.02 |
| IMAGE−I | 700 | 700 | 1.03 |
| UK | 721 | 5081 | 1.13 |
| Brazil children | 259 | 259 | 1.1 |
| Spain children | 430 | 639 | 1.12 |
| Denmark children | 6462 | 5879 | 1.13 |
| Summary | 15591 | 29722 | 1.1 |

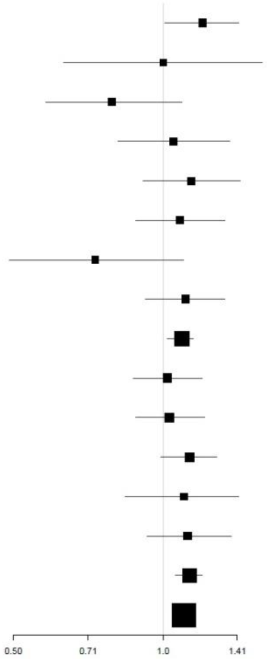


D.

**rs1920644**

| Study | Cases | Controls | OR |
|---|---|---|---|
| Germany adults | 552 | 1062 | 1.09 |
| The Netherlands | 124 | 112 | 1.37 |
| Norway I | 305 | 208 | 0.84 |
| Norway II | 390 | 354 | 1.14 |
| Brazil adults | 417 | 463 | 1.26 |
| Spain I | 418 | 428 | 1.04 |
| Spain II | 126 | 126 | 1.38 |
| Spain III | 359 | 1414 | 1.09 |
| Denmark adults | 3841 | 11707 | 1.06 |
| PUWMa, USA | 563 | 563 | 1.01 |
| Canada | 109 | 109 | 1.36 |
| Germany children | 487 | 1290 | 1.26 |
| IMAGE−I | 700 | 700 | 1.15 |
| IMAGE−II | 624 | 1755 | 0.92 |
| CHOP, USA | 262 | 262 | 1.07 |
| UK | 721 | 5081 | 1.12 |
| Brazil children | 259 | 259 | 1.03 |
| Spain children | 430 | 639 | 1.26 |
| Denmark children | 6462 | 5879 | 1.07 |
| **Summary** | **17149** | **32411** | **1.09** |

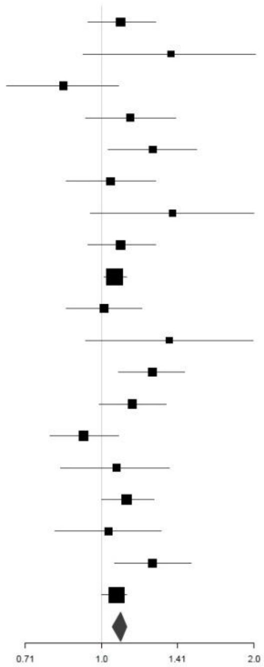

**Supplementary Table 1. Sample description, quality control filters and more details for each new cohort included in the present study [excel file].** Information for each new GWAS included in the present study is provided. For the other GWAS (CHOP (USA), IMAGE-I (Europe), PUWMa (USA), Canada (Toronto), UK (Cardiff), Germany, and IMAGE-II (Europe and USA)) extensive information is available in Demontis and Walters et al.[31]

**Supplementary Table 2. Genes surviving Bonferroni correction in the gene-based analysis of (A) persistent ADHD and (B) ADHD in childhood (with P<2.77E-06 and P<2.75E-06, respectively).**

**A.**

| Gene | Chr | Start | Stop | N SNPs* | N PARAM** | Z-STAT | P-value |
|---|---|---|---|---|---|---|---|
| *ST3GAL3* | 1 | 44153204 | 44416837 | 535 | 37 | 4.8 | 8.72E-07 |
| *FRAT1* | 10 | 99059022 | 99101672 | 150 | 16 | 4.6 | 1.98E-06 |
| *FRAT2* | 10 | 99072254 | 99114458 | 124 | 17 | 4.6 | 1.86E-06 |
| *CGB1* | 19 | 49518826 | 49560191 | 98 | 15 | 4.6 | 1.82E-06 |
| *RNF225* | 19 | 58887457 | 58928446 | 122 | 15 | 4.6 | 2.38E-06 |
| *ZNF584* | 19 | 58899245 | 58949694 | 142 | 12 | 4.7 | 1.47E-06 |

**B.**

| Gene | Chr | Start | Stop | N SNPs* | N PARAM** | Z-STAT | P-value |
|---|---|---|---|---|---|---|---|
| *FEZF1* | 7 | 121921373 | 121971173 | 92 | 15 | 4.9 | 5.42E-07 |

Chr: chromosome
* N SNPs: number of single nucleotide polymorphisms in the genes.
** N PARAM: number of relevant parameters used in the model by MAGMA software [37].

**Supplementary Table 3. Top hits (P<5.00E-06) in the GWAS meta-analysis of (A) persistent ADHD and (B) ADHD in childhood.** The location (chromosome (Chr) and base position (BP)), effect allele and its frequency, odds ratio (OR) of the effect allele with 95% confidence interval (CI 95%), association P-values and direction of effect for each cohort are shown for each index variant ID (SNP).

**A.**

| Chr | BP | SNP | Effect allele | Freq effect allele | OR | CI 95% | P-value |
|---|---|---|---|---|---|---|---|
| 8 | 88413971 | rs3923931 | A | 0.11 | 1.20 | 1.12-1.28 | 1.69E-07 |
| 9 | 119656429 | rs4836899 | T | 0.52 | 1.12 | 1.07-1.17 | 2.76E-07 |
| 9 | 17091464 | rs10962864 | T | 0.16 | 1.16 | 1.09-1.23 | 5.07E-07 |
| 14 | 29754596 | rs28768443 | A | 0.19 | 0.87 | 0.82-0.92 | 6.22E-07 |
| 2 | 155361952 | rs13019774 | T | 0.39 | 1.12 | 1.07-1.17 | 1.20E-06 |
| 10 | 101904937 | rs35835615 | CA | 0.91 | 0.83 | 0.77-0.90 | 1.27E-06 |
| 9 | 86582923 | rs296884 | T | 0.26 | 0.89 | 0.84-0.93 | 1.40E-06 |
| 18 | 22644275 | rs55654452 | G | 0.56 | 1.11 | 1.06-1.16 | 1.65E-06 |
| 16 | 86394764 | rs113773212 | A | 0.95 | 0.78 | 0.71-0.86 | 1.66E-06 |
| 1 | 159675717 | rs876538 | T | 0.21 | 0.88 | 0.83-0.92 | 1.70E-06 |
| 1 | 30088755 | rs9661242 | A | 0.52 | 0.90 | 0.86-0.94 | 1.94E-06 |
| 15 | 81817631 | rs74030106 | A | 0.95 | 0.78 | 0.70-0.86 | 2.27E-06 |
| 10 | 2782803 | rs1537617 | A | 0.84 | 1.17 | 1.10-1.25 | 2.44E-06 |
| 1 | 215652459 | rs6540899 | A | 0.12 | 1.21 | 1.12-1.31 | 2.47E-06 |
| 9 | 17044971 | rs393488 | A | 0.47 | 0.90 | 0.86-0.94 | 2.63E-06 |
| 9 | 17263748 | rs145190003 | T | 0.74 | 0.89 | 0.85-0.94 | 2.99E-06 |
| 13 | 97857492 | rs57422381 | A | 0.93 | 0.82 | 0.75-0.89 | 3.17E-06 |
| 3 | 8165502 | rs13088735 | T | 0.02 | 1.48 | 1.25-1.75 | 3.37E-06 |
| 6 | 92066647 | rs35854754 | CT | 0.94 | 1.27 | 1.15-1.40 | 3.39E-06 |
| 3 | 97917645 | rs75311156 | T | 0.18 | 0.86 | 0.81-0.92 | 3.71E-06 |
| 19 | 58926345 | rs35782676 | T | 0.77 | 1.13 | 1.07-1.19 | 3.73E-06 |
| 4 | 188166018 | rs34846424 | T | 0.36 | 1.11 | 1.06-1.16 | 3.77E-06 |
| 21 | 30109998 | rs2832008 | T | 0.86 | 0.87 | 0.81-0.92 | 3.96E-06 |
| 4 | 56658778 | rs895614 | A | 0.29 | 1.13 | 1.07-1.19 | 4.00E-06 |
| 9 | 17162413 | rs6475111 | T | 0.26 | 1.13 | 1.07-1.18 | 4.56E-06 |

**B.**

| Chr | BP | SNP | Effect allele | Freq Effect allele | OR | CI 95% | P-value |
|---|---|---|---|---|---|---|---|
| 16 | 8508591 | rs55686778 | T | 0.19 | 1.15 | 1.09-1.21 | 1.67E-07 |
| 1 | 175322306 | rs191624305 | T | 0.90 | 1.21 | 1.13-1.30 | 2.47E-07 |
| 22 | 17487476 | rs138596453 | A | 0.02 | 0.67 | 0.57-0.79 | 6.33E-07 |
| 15 | 47806012 | rs1610098 | T | 0.63 | 0.90 | 0.87-0.94 | 7.62E-07 |
| 16 | 73024276 | rs1858800 | T | 0.35 | 1.12 | 1.07-1.17 | 1.14E-06 |
| 4 | 140729346 | rs795989 | A | 0.68 | 1.11 | 1.07-1.17 | 1.53E-06 |
| 10 | 8850735 | rs11255938 | T | 0.46 | 0.91 | 0.87-0.94 | 2.16E-06 |
| 2 | 47505000 | rs79221481 | T | 0.03 | 1.39 | 1.21-1.60 | 2.60E-06 |
| 8 | 13664469 | rs536749404 | T | 0.14 | 1.17 | 1.10-1.25 | 2.66E-06 |
| 6 | 117391879 | rs526318 | T | 0.54 | 1.10 | 1.06-1.15 | 2.76E-06 |

**Supplementary Table 4. Pathway analyses (A) on persistent ADHD (B) on ADHD in childhood and (C) on ADHD across the lifespan. [excel file]**

**Supplementary Table 5. Sign test results for (A) variants associated with ADHD in childhood in persistent ADHD and (B) variants associated with persistent ADHD in ADHD in childhood.**

**A.**

| P-value threshold | All SNPs (N) | SNPs showing the same sign (N) | Proportion SNPs with the same sign (%) | 95% CI | P-value* |
|---|---|---|---|---|---|
| 5.00E-04 | 727 | 441 | 60.66 | 0.57-0.64 | <2.22E-16 |
| 1.00E-04 | 200 | 127 | 63.50 | 0.56-0.70 | 1.80E-04 |
| 5.00E-05 | 118 | 78 | 66.10 | 0.57-0.74 | 6.60E-04 |
| 1.00E-05 | 30 | 23 | 76.67 | 0.57-0.89 | 6.17E-03 |
| 5.00E-06 | 17 | 12 | 70.59 | 0.44-0.89 | 0.15 |
| 1.00E-06 | 4 | 3 | 75.00 | 0.22-0.99 | 0.62 |
| 5.00E-07 | 2 | 1 | 50.00 | 0.09-0.91 | 1 |

**B.**

| P-value threshold | All SNPs (N) | SNPs showing the same sign (N) | Proportion SNPs with the same sign (%) | 95% CI | P-value* |
|---|---|---|---|---|---|
| 5.00E-04 | 743 | 442 | 59.49 | 0.56-0.63 | <2.22E-16 |
| 1.00E-04 | 198 | 128 | 64.65 | 0.58-0.71 | 5.00E-05 |
| 5.00E-05 | 113 | 72 | 63.72 | 0.54-0.72 | 4.47E-03 |
| 1.00E-05 | 30 | 19 | 63.33 | 0.44-0.79 | 0.20 |
| 5.00E-06 | 21 | 12 | 57.14 | 0.34-0.77 | 0.66 |
| 1.00E-06 | 4 | 3 | 75.00 | 0.22-0.99 | 0.62 |
| 5.00E-07 | 2 | 1 | 50.00 | 0.09-0.91 | 1 |

CI: confidence interval
*P-values <2.22E-16 in R were not provided

**Supplementary Table 6. Quintile results for association of PRS for ADHD in childhood with persistent ADHD. Meta-analysis P-values at threshold P=0.4 in the discovery analysis (GWAS meta-analysis of ADHD in childhood) is shown.**

| Quintile | OR | 95% CI* | P-value |
|---|---|---|---|
| 1 | 0.92 | 0.88-0.96 | 2.35E-04 |
| 2 | 0.96 | 0.92-1.00 | 7.27E-02 |
| 3 | 1 | 1 | NA |
| 4 | 1.06 | 1.02-1.11 | 7.69E-03 |
| 5 | 1.13 | 1.08-1.18 | 2.36E-08 |

CI: confidence interval

**Supplementary Table 7. Results of the BUHMBOX analysis.** BUMHMBOX analysis results, testing for subgroup heterogeneity with respect to persistent ADHD among individuals with ADHD in childhood, are shown for 62 variants with P<5.00E-05 (indicated with *) and for 710 variants with P<1.00E-03 (indicated with **) in the GWAS meta-analysis of persistent ADHD.

| **Study** | **Neff** | **Z-score*** | **P-value*** | **Z-score**** | **P-value**** |
|---|---|---|---|---|---|
| Brazil | 259.00 | -0.0170 | 0.51 | 1.6715 | 0.05 |
| Denmark | 6156.73 | -1.8068 | 0.97 | -0.9916 | 0.84 |
| Germany | 707.07 | -0.8294 | 0.80 | 1.5340 | 0.06 |
| IMAGE-I | 700.00 | -0.6192 | 0.73 | 0.8285 | 0.20 |
| IMAGE-II | 920.66 | -0.4724 | 0.68 | -1.2291 | 0.89 |
| UK | 1262.81 | 0.9749 | 0.16 | -1.1632 | 0.88 |
| Canada | 109.00 | 1.4409 | 0.07 | -0.1835 | 0.57 |
| Spain | 514.07 | 1.1129 | 0.13 | -0.4641 | 0.68 |
| CHOP, USA | 262.00 | 0.4837 | 0.31 | -1.2575 | 0.90 |
| PUWMa, USA | 563.00 | -0.6192 | 0.73 | -2.1844 | 0.99 |
| Overall | | -1.1844 | 0.88 | -1.5815 | 0.94 |

Neff: effective sample size

**Supplementary Table 8. Comparison of genome-wide significant hits of the GWAS meta-analysis of ADHD across the lifespan with genome-wide significant hits of the previous GWAS meta-analysis of ADHD by Demontis and Walters et al., 2019[31].**

(A) Results of the genome-wide significant variants of the GWAS meta-analysis of ADHD across the lifespan in the GWAS-MA by Demontis and Walters et al.[31]; (B) Results of the genome-wide significant variants reported by Demontis and Walters et al.[31] in the GWAS meta-analysis of ADHD across the lifespan; (C) Results of the genome-wide significant genes reported by Demontis and Walters et al. [31] in the gene-based analysis of the GWAS meta-analysis of ADHD across the lifespan.

**A.**

| | | | | **Present study** | | | **Demontis and Walters et al., 2019** | | |
|---|---|---|---|---|---|---|---|---|---|
| **Chr** | **BP** | **SNP** | **Effect allele** | **OR** | **CI 95%** | **P-value** | **OR** | **CI 95%** | **P-value** |
| 6 | 159384224 | rs183882582 | T | 1.43 | 1.26-1.60 | 1.57E-08 | 1.18 | 1.07-1.30 | 1.24E-03 |
| 7 | 121955328 | rs3958046 | T | 1.09 | 1.06-1.10 | 2.28E-08 | 1.07 | 1.04-1.10 | 5.09E-07 |
| 4 | 31151465 | rs200721207 | T | 1.10 | 1.06-1.13 | 3.56E-08 | 1.09 | 1.06-1.12 | 1.15E-08 |
| 3 | 160313354 | rs1920644 | T | 1.09 | 1.05-1.12 | 4.74E-08 | 1.05 | 1.03-1.08 | 1.56E-04 |

**B.**

| | | | | **Demontis and Walters et al., 2019** | | **Present study** | | |
|---|---|---|---|---|---|---|---|---|
| **Chr** | **BP** | **SNP** | **Effect allele** | **OR** | **P-value** | **OR** | **CI 95%** | **P-value** |
| 1 | 44184192 | rs11420276 | G | 1.11 | 2.14E-13 | 1.09 | 1.04-1.13 | 2.53E-04 |
| 1 | 96602440 | rs1222063 | A | 1.10 | 3.07E-08 | 1.07 | 1.01-1.14 | 2.27E-02 |
| 2 | 215181889 | rs9677504 | A | 1.12 | 1.39E-08 | 1.08 | 1.03-1.13 | 8.59E-04 |
| 3 | 20669071 | rs4858241 | T | 1.08 | 1.74E-08 | 1.08 | 1.05-1.12 | 4.79E-07 |
| 4 | 31151456 | rs28411770 | T | 1.09 | 1.15E-08 | 1.11 | 1.06-1.17 | 7.62E-06 |
| 5 | 87854395 | rs4916723 | A | 0.93 | 1.58E-08 | 0.95 | 0.92-0.97 | 2.89E-04 |
| 7 | 114086133 | rs5886709 | G | 1.08 | 1.66E-08 | 1.05 | 1.02-1.08 | 1.17E-03 |
| 8 | 34352610 | rs74760947 | A | 0.84 | 1.35E-08 | 0.86 | 0.80-0.92 | 1.15E-05 |
| 10 | 106747354 | rs11591402 | A | 0.91 | 1.34E-08 | 0.94 | 0.90-0.97 | 2.12E-04 |
| 12 | 89760744 | rs1427829 | A | 1.08 | 1.82E-09 | 1.08 | 1.05-1.11 | 5.64E-07 |
| 15 | 47754018 | rs281324 | T | 0.93 | 2.68E-08 | 0.93 | 0.90-0.96 | 7.22E-07 |
| 16 | 72578131 | rs212178 | A | 0.89 | 7.68E-09 | 0.91 | 0.87-0.96 | 1.36E-04 |

Chr: chromosome
BP: base pair
OR: odds ratio
CI: confidence interval

**C.**

| Gene | Chr | Start | Stop | Demontis and Walters et al., 2019<br>P-value | Present study<br>P-value |
|---|---|---|---|---|---|
| *ST3GAL3* | 1 | 44173204 | 44396837 | 7.38E-12 | 3.58E-08 |
| *KDM4A* | 1 | 44115797 | 44171189 | 2.15E-11 | 4.34E-07 |
| *PTPRF* | 1 | 43991708 | 44089343 | 5.68E-10 | 2.78E-05 |
| *SZT2* | 1 | 43855556 | 43919918 | 8.47E-09 | 9.27E-04 |
| *TIE1* | 1 | 43766566 | 43788781 | 2.01E-08 | 2.29E-03 |
| *MPL* | 1 | 43803475 | 43820135 | 3.33E-08 | 2.25E-03 |
| *CDC20* | 1 | 43824626 | 43828874 | 6.34E-08 | 2.16E-03 |
| *HYI* | 1 | 43916674 | 43919938 | 3.28E-07 | 1.01E-03 |
| *SLC6A9* | 1 | 44462155 | 44497171 | 7.58E-07 | 3.23E-04 |
| *ELOVL1* | 1 | 43829068 | 43833745 | 1.26E-06 | 1.82E-03 |
| *CCDC24* | 1 | 44457280 | 44462200 | 2.12E-06 | 5.97E-05 |
| *MANBA* | 4 | 103552643 | 103682151 | 6.00E-08 | 1.13E-05 |
| *MEF2C* | 5 | 88014058 | 88199922 | 3.19E-08 | 1.55E-05 |
| *FOXP2* | 7 | 113726365 | 114333827 | 5.50E-07 | 4.48E-05 |
| *SORCS3* | 10 | 106400859 | 107024993 | 2.18E-09 | 6.00E-05 |
| *CUBN* | 10 | 16865965 | 17171816 | 1.59E-07 | 3.28E-04 |
| *PIDD1* | 11 | 799179 | 809872 | 5.30E-07 | 1.82E-04 |
| *DUSP6* | 12 | 89741837 | 89746296 | 2.24E-09 | 3.51E-08 |
| *SEMA6D* | 15 | 47476403 | 48066420 | 2.63E-10 | 7.24E-08 |
| *CDH8* | 16 | 61681169 | 62070939 | 4.67E-08 | 1.71E-04 |

**Supplementary Table 9. Bayesian credible sets for the four loci identified in the meta-analysis of GWAS on ADHD across the lifespan, annotation and eQTL information. [excel file]**

**Supplementary Table 10. Summary-data-based Mendelian Randomization (SMR) results in blood and brain.** Significant association after Bonferroni correction is highlighted in grey. $P_{HEIDI}$ indicates the P value for the Heterogeneity in dependent instruments (HEIDI) to distinguish pleiotropy from linkage. **[excel file]**

**Supplementary Table 11. Genetic correlations between ADHD datasets**

| Tested datasets* | rg (%) | SE | P-value** |
|---|---|---|---|
| aADHD vs PGC+iPSYCH (EUR) | 85.06 | 0.04 | 5.49E-99 |
| cADHD vs PGC+iPSYCH (EUR) | 98.97 | 0.03 | 5.02E-273 |
| a+cADHD- PGC+iPSYCH (EUR) | 98.47 | 0.01 | <2.23E-308 |
| aADHD vs EAGLE | 65.27 | 0.20 | 1.10E-03 |
| cADHD vs EAGLE | 97.78 | 0.21 | 2.76E-06 |
| a+cADHD- EAGLE | 87.06 | 0.19 | 4.80E-06 |
| aADHD vs 23andMe | 74.86 | 0.05 | 2.49E-45 |
| cADHD vs 23andMe | 63.25 | 0.05 | 1.39E-42 |
| a+cADHD- 23andMe | 72.16 | 0.04 | 4.86E-88 |

*aADHD: adult ADHD; cADHD: children ADHD; a+cADHD: ADHD across lifetime, EUR: European
**P-values <2.23E-308 in Python were not provided

**Supplementary Table 12. Genetic correlation of ADHD in childhood, persistent ADHD in adults and ADHD across the lifespan with other traits [excel file].** Traits showing significant genetic correlation (rg) with persistent ADHD in adults and with ADHD in childhood after Bonferroni correction are highlighted in bold.
